# Rookognise: Acoustic detection and identification of individual rooks in field recordings using multi-task neural networks

**DOI:** 10.1101/2022.02.19.481011

**Authors:** Killian Martin, Olivier Adam, Nicolas Obin, Valérie Dufour

**Affiliations:** PRC, UMR 7247, Ethologie Cognitive et Sociale, INRAE-CNRS-IFCE-Université de Tours, Strasbourg, France; Institut Jean Le Rond d’Alembert, UMR 7190, Sorbonne Université-CNRS, F-75005 Paris, France; Institut des Neurosciences Paris-Saclay, UMR 9197, CNRS-Université Paris Sud, Orsay, France; STMS Lab, IRCAM, CNRS, Sorbonne Université, Paris, France

**Keywords:** deep learning, bird recognition, outdoor setting, vocalisation-independent, corvid, multi-channel audio

## Abstract

Individual-level monitoring is essential in many behavioural and bioacoustics studies. Collecting and annotating those data is costly in terms of human effort, but necessary prior to conducting analysis. In particular, many studies on bird vocalisations also involve manipulating the animals or human presence during observations, which may bias vocal production. Autonomous recording units can be used to collect large amounts of data without human supervision, largely removing those sources of bias. Deep learning can further facilitate the annotation of large amounts of data, for instance to detect vocalisations, identify the species, or recognise the vocalisation types in recordings. Acoustic individual identification, however, has so far largely remained limited to a single vocalisation type for a given species. This has limited the use of those techniques for automated data collection on raw recordings, where many individuals can produce vocalisations of varying complexity, potentially overlapping one another, with the additional presence of unknown and varying background noise. This paper aims at bridging this gap by developing a system to identify individual animals in those difficult conditions. Our system leverages a combination of multi-scale information integration, multi-channel audio and multi-task learning. The multi-task learning paradigm is based the overall task into four sub-tasks, three of which are auxiliary tasks: the detection and segmentation of vocalisations against other noises, the classification of individuals vocalising at any point during a sample, and the sexing of detected vocalisations. The fourth task is the overall identification of individuals. To test our approach, we recorded a captive group of rooks, a Eurasian social corvid with a diverse vocal repertoire. We used a multi-microphone array and collected a large scale dataset of time-stamped and identified vocalisations recorded, and found the system to work reliably for the defined tasks. To our knowledge, the system is the first to acoustically identify individuals regardless of the vocalisation produced. Our system can readily assist data collection and individual monitoring of groups of animals in both outdoor and indoor settings, even across long periods of time, and regardless of a species’ vocal complexity. All data and code used in this article is available online.

## 1 Introduction

Deep learning systems have become invaluable tools in the ecological sciences in recent years. They allow the automation of many monitoring tasks, enabling much wider spatio-temporal coverage than previous methods (Christin et al., 2019; Weinstein, 2019). Deep learning system have been applied to tasks as diverse as detecting species in complex environments (e.g Conrady et al., 2022; Dufourq et al., 2022; Fu et al., 2022; She et al., 2022; van Klink et al., 2022), censusing populations (Adi et al., 2010), surveying breeding success of potentially endangered species (Teixeira et al., 2022), or tracking invasive species both for plants and animals (Campos et al., 2022; Li et al., 2021; Takimoto et al., 2021). Beyond the species level, observing individual animals remains a challenge for researchers (Ferreira et al., 2020) despite its essential role in ecological and behavioural studies (Clutton-Brock and Sheldon, 2010; Terry et al., 2005). Extensive research has evidenced the possibility of identifying individual animals by their vocalisations (see e.g. Beecher, 1989; Bradbury and Vehrencamp, 1998; Linhart et al., 2019), opening the possibility of applying passive acoustic monitoring techniques (Schneider et al., 2019) to individual monitoring. Bird vocalisations have been studied in the ecological and evolutionary sciences due to their pervasive importance to the life history of birds (see e.g. Catchpole and Slater (2008) and Marler and Slabbekoorn (2004) for reviews), but also because of their potential analogies with human language (e.g. Sainburg et al., 2019). Current research questions investigate, for example, the interplay between vocal production and sociality in the form of conversations, dialects and vocal signatures in many bird species.

The vocal signature is the set of bioacoustic features that are individually distinctive within a given call type. Such signatures have been found throughout the animal kingdom (Jansen et al., 2012; Kershenbaum et al., 2013; Linhart et al., 2019; McCordic et al., 2016). However, most studies in vocal signature have been limited to a single vocalisation type (e.g. Benti et al., 2019; Boeckle et al., 2012, 2018; Laiolo et al., 2000; Mates et al., 2015; Stowell et al., 2016; Stowell et al., 2019a; Yorzinski et al., 2006). When encompassing multiple different vocalisation types, the concept of vocal signature becomes a lot more nebulous, as different vocalisation types can bear different signatures (Elie and Theunissen, 2018; Keenan et al., 2020). This plurality of signatures obviously complicates the identification of individuals in species with a rich repertoire, at least with classical methods (Catchpole and Slater, 2008; Elie and Theunissen, 2018). Despite this, there have been a few cases of successful individual identification with multiple vocalisation types (Cheng et al., 2012; Fox et al., 2008; Kirschel et al., 2009), most successfully with pre-deep learning era neural networks. Still, these approaches were based on supervised algorithms trained on vocalisations manually extracted from the recordings, limiting their applicability to field recordings (Kershenbaum et al., 2014; Thompson et al., 1994).

Neural networks appear as the most appropriate tool to acoustically classify birds down to the individual level. Recent breakthroughs in general machine learning and specifically deep learning (Stowell, 2022) have brought state-of-the-art results just in bird vocalisation detection (Fanioudakis and Potamitis, 2017; Grill and Schlüter, 2017; Kong et al., 2017; Liaqat et al., 2018; Lostanlen et al., 2019; Sevilla and Glotin, 2017; Stowell et al., 2019b), species classification (Kahl et al., 2019; Kahl et al., 2021), and segmentation of song with classification of vocalisation types (Cohen et al., 2022). But the deep learning community has so far only rarely investigated the problem of individual identification. One possible explanation is that *in situ* recording remains challenging, in particular recording individual birds (Folliot et al., 2022; Stowell et al., 2019a), due to interferences from other sound sources, including conspecifics, and simple wariness preventing the observer from approaching enough to obtain many recordings of good quality. As such, many studies relying on individual recordings simply record the bird alone in a separate chamber (e.g. Sainburg et al., 2019); however, this experimental setting deprives birds of their natural and social environment and can bias vocal production. In addition, acquiring manually individually annotated data is time consuming and generally costly (Stowell et al., 2019a). Autonomous recording units (ARUs) can be used to record audio in a more ecologically valid environment, i.e. outdoor, in a social group, and without the need for human presence. They have been increasingly used in ecological surveys, since they allow wide coverage with little human effort (Blumstein et al., 2011; Darras et al., 2019; Fristrup and Mennitt, 2012; Potamitis et al., 2014; Schneider et al., 2019; Shonfield and Bayne, 2017).

Deep learning is adequately suited to process the vast amount of complex data that can be collected with ARUs thanks to the representational power offered by these networks. Furthermore, the network’s architecture can be adapted to treat the various tasks at (in our case, detection of vocalisation and individualisation). For instance, multiple tasks can be tackled using a multi-task learning approach, i.e. by learning the tasks jointly within a single network. This approach enables the network to learn a shared representation, which has been suggested to effectively bias the learning process towards representations that benefit all tasks (Caruana, 1997; Liebel and Körner, 2018; Ruder, 2017). This shared representation can not only enhance training, but also the generalisation ability of the network. This is usually implemented by sharing early parts of the network, before splitting into task-specific heads. For instance, one head may learn to detect objects over the background, and another may learn to classify the detected objects (Morfi and Stowell, 2018; Pankajakshan et al., 2019). In our approach, we define four such sub-tasks: the *detection* sub-task whose objective is to detect for each audio frame whether a bird is vocalising or not (versus any other sound), the *sexing* sub-task whose objective is to further classify whether the individual producing the vocalisation is a male or a female, the *presence* sub-task whose objective is to identity the presence of a vocalisation of a particular individual within a large audio clip regardless to its precise time position, and the *identification* sub-task, which returns time-stamped vocalisations with the identity of the individual producing each vocalisation.

We then tested the performance of the system on rooks (*Corvus frugilegus*), held in a social group and in an outdoor setting. Rooks are a European social corvid with a rich vocal repertoire (Røskaft and Espmark, 1982). Corvids are known for producing a large number of different vocalisations with a chaotic structure and a wide frequency range (Brown, 1985; Marzluff and Angell, 2005; Røskaft and Espmark, 1982), particularly difficult to parametrise using classical acoustic measures (Fagerlund and Härmä, 2005; Fletcher, 2000). Furthermore, individuals only share some of their repertoire with their conspecifics (Boeckle et al., 2012; Kondo et al., 2010; Mates et al., 2015), further limiting the use of classical methods for individual identification. Rooks, in particular, may possess an even richer repertoire than previously thought, as they produce series of vocalisations similar to the song of many other oscines (Coombs, 1960), that have never been described in detail. While further description is beyond the scope of this paper, rooks appear to be of particular interest to develop a network capable of vocalisation-independent individual identification.

To summarise, in this paper we present the following contributions, to overcome the afore-mentioned limitations:

1. We present the first system able to acoustically identify individual birds whatever their vocalisation types, which detects vocalisations from raw recordings and attributes them to the individual emitter. Furthermore, the system was developed using data sampled in an outdoors aviary and was trained to learn its tasks in an environment with multiple sources of noise. The system is trained in a fully supervised manner, on a dataset of identified rook vocalisations. The code for this system is available online.
2. The architecture is based on a multi-task learning approach. This architecture is used in order to learn information useful for multiple related tasks, and to increase the generalisation ability of the system compared to multiple single-task networks. The multi-task approach exploits information from a shared representation learned by all tasks, followed by task-specific heads. This formulation also lets us fuse the predictions of several sub-tasks together as an attention-like mechanism (e.g. where the detection head returns a low score, there is no need for the identification head to learn to identify an individual). Furthermore, a multi-scale module is added to allow the network to exploit information from very local cues in the spectrogram up to vocalisation-wide patterns to find individually-distinctive information.
3. The proposed multi-task network operates on multi-channel audio signals, as recorded from a multi-microphone array. This allows the system to exploit spatial information to enhance the detection of rook vocalisations as well as the subsequent tasks. The multi-channel recording also adds redundancy in the case of degraded, noisy or weak signals (e.g. saturation due to a bird vocalising close to the microphone, or strong non-vocal noises). To our knowledge, this is the first implementation of a neural network that exploits spatial information in the domain acoustic individual identification.
4. We collected the first large scale annotated dataset of rook vocalisations. This dataset includes vocalisations from 15 individual rooks collected over the course of one year, with each vocalisation time-stamped and attributed to the individual producing it. The data collection involved a multi-microphone array spread throughout the aviary to record each bird from as close as possible. As such, the recordings are composed of multi-channel audio, unlike most approaches in the literature. The entire annotated dataset is freely available online under a Creative Common (CC/BY 4.0) Licence^2^.

## 2 Material and methods

### 2.1 Data collection

We recorded the vocalisations produced by a captive colony of 15 adult rooks (8 males, 7 females) housed in an outdoor aviary in Strasbourg, France. All birds were identified by coloured leg rings and had been housed together since they had been caught as fledglings, except one adult male added shortly before the start of the recordings. We recorded the group using a multi-microphone array constituted of up to three ARUs, each with two microphones on 3 m cables (Song Meter 4 and SMM-A2 microphones, Wildlife Acoustics, USA). The ARUs were programmed to record for several consecutive hours each morning from January 2020 onwards. The microphones were spaced across the aviary so the distance between microphone and bird was at most approximately 10 m. The resulting multi-channel recordings were digitised at 48 kHz with a 16-bit resolution. A schematic layout of the aviary, with microphone placements and the usual spots the birds stayed at during observations, can be found in the supplementary material (Fig. A.1).

Among the daily recordings, eleven long sessions and a number of shorter recordings were conducted in the presence of an expert observer (KM). During these sessions, a custom Python program allowed the observer to note the identity of the bird that emitted each vocalisation (thanks to their leg rings). The program also recorded the time of each annotation, which later allowed synchronisation with the audio recordings. These on-site annotations were completed with video recordings also synchronisable to both the annotation and audio files. These videos were used to help resolve potential ambiguities (e.g. to identify which one of two birds vocalised if both were close to each other). These synchronised files, obtained throughout 2020 and 2021 (see Fig. A.2 for a timeline) were used to build an annotated dataset by the same expert observer with the AudioSculpt software (Bogaards et al., 2004). Each vocalisation was annotated with the start time, end time, emitter identity, and whether it was part of a sequence. Due to the multi-microphone array and sound propagation speed, the start and end times were defined as the time when a vocalisation was perceived on any audio channel for the first and last time, respectively. A vocalisation was defined as either an uninterrupted sound or a short succession of small, similar elements that were always produced together. Despite both notes and video assistance, ambiguities sometimes remained regarding the identity of emitters. When this was the case, the emitter was noted as either ”Unknown” (single emitter could not be identified), or ”Multiple” (multiple emitters vocalised simultaneously). The final annotated dataset included 17,662 vocalisations in 17.4h of audio recordings. The vocalisations were not equally distributed among individuals; one male in particular vocalised much more often than all the other birds (Fig. 1). Rooks, like many birds, vocalise following two general modes, calls and song-like sequences. Since our objective is to identify individuals regardless of what vocalisations they produce, we do not separate the two classes in this study.

**Figure 1:**
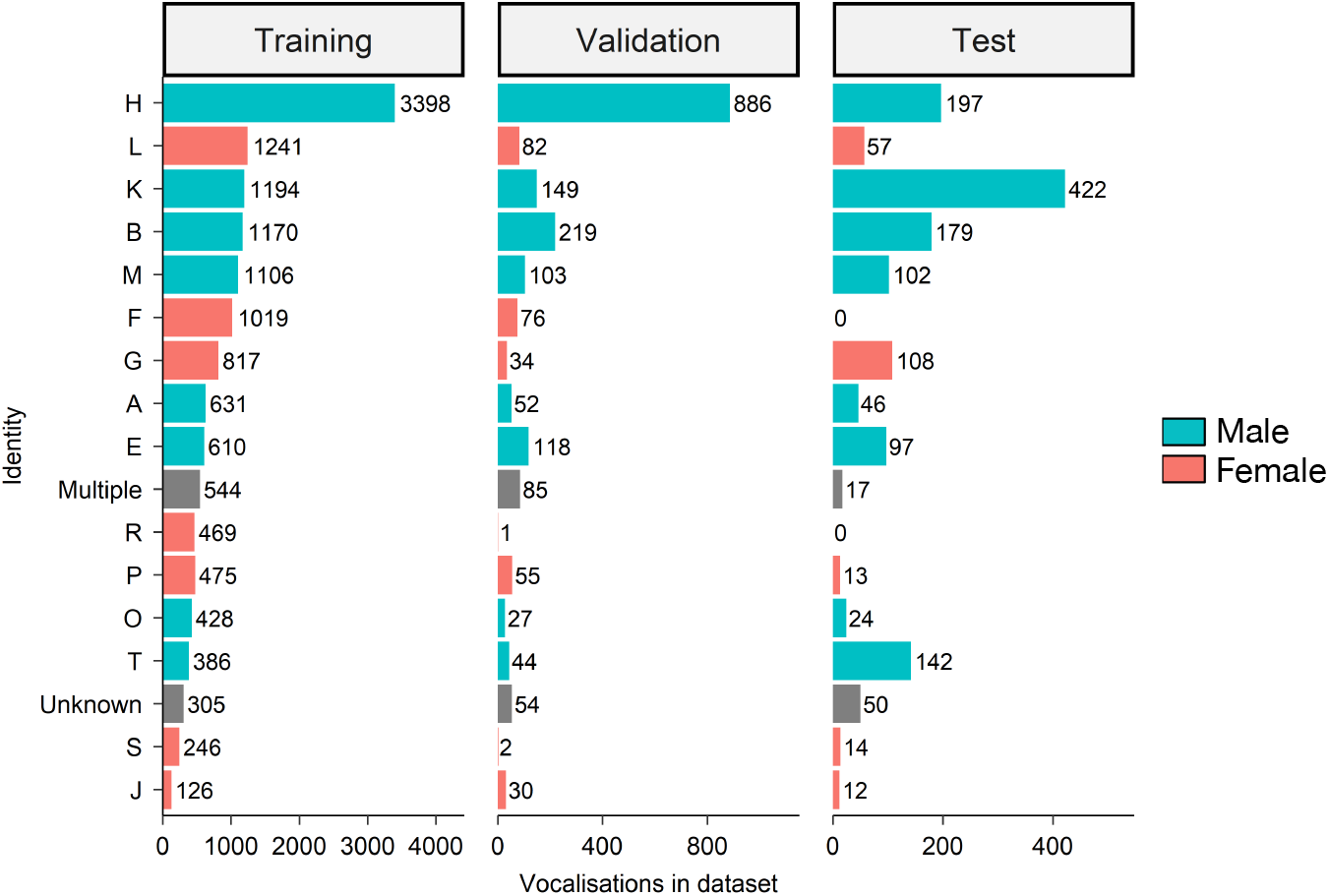
Number of vocalisations per bird and dataset, in decreasing order based on the training dataset. The numbers next to the bars represent the exact number of vocalisations produced by each bird in this dataset. Bar colours represent the sex of the individual. Multiple and Unknown represent cases where vocalisations could not be attributed to an individual bird.

All EU ethical guidelines (Directive 2010/63/EU) were followed for the care of the rooks throughout the study. No experimental procedure was necessary for the data collection.

### 2.2 Audio representation

From the multi-channel raw audio recordings, we extracted Mel-spectrograms corresponding to 10 s of audio, using all microphones available (up to 6). The length of the clips was based on Grill and Schlüter (2017), itself based on the lengths of most clips in the Bird Audio Detection Challenge dataset (Stowell et al., 2019b). This length was the largest value that fit in computer memory with the batch size chosen for the networks. We extracted 10 s clips, applied a pre-emphasis filter with the following equation: *y*[*t*] = *x*[*t*] *− x*[*t −* 1], where *x*[*t*] is the original audio at time *t* and *y*[*t*] is the corresponding filter output. From the output of this filter, we computed the Short Time Fourier Transform of each clip using a 50 ms Hamming window with 75% overlap, then extracted the squared magnitude of the output. The spectrograms were then downscaled to 80 Mel-scale frequency coefficients to emphasise lower frequencies. We chose these parameters empirically based on the values that produced spectrograms with good time and frequency resolution during the annotations. This resulted in Mel-spectrograms with dimensions (*F, T,C*), with *F* the number of Mel-coefficients, *T* the number of spectrogram time frames (800 in our implementation), and *C* the number of microphones (6 in our implementation). Smaller clips were zero-padded at the end of each dimension.

### 2.3 Network architecture

#### Shared front-end

The front-end (illustrated in Fig. 2B) encodes information from the input Mel-spectrograms through successive convolution and pooling operations. The first layer is a Per-Channel Energy Normalisation layer (PCEN, Wang et al., 2017) implemented so that all parameters of the PCEN operation are learned during the training process, with each parameter encoded as a vector of size 80, so that each frequency band has its own value for each parameter of the PCEN equation. The rest of its architecture is based on the sparrow network proposed by Grill and Schlüter (2017), variants of which were used in Schlüter (2018) and Kahl et al. (2021) to classify bird species with excellent results. Following Grill and Schlüter (2017), we standardise each frequency band and audio channel separately with a Batch Normalisation layer between the PCEN layer and the rest of the front-end. Formally, the front-end corresponds to the following equation:

**Figure 2:**
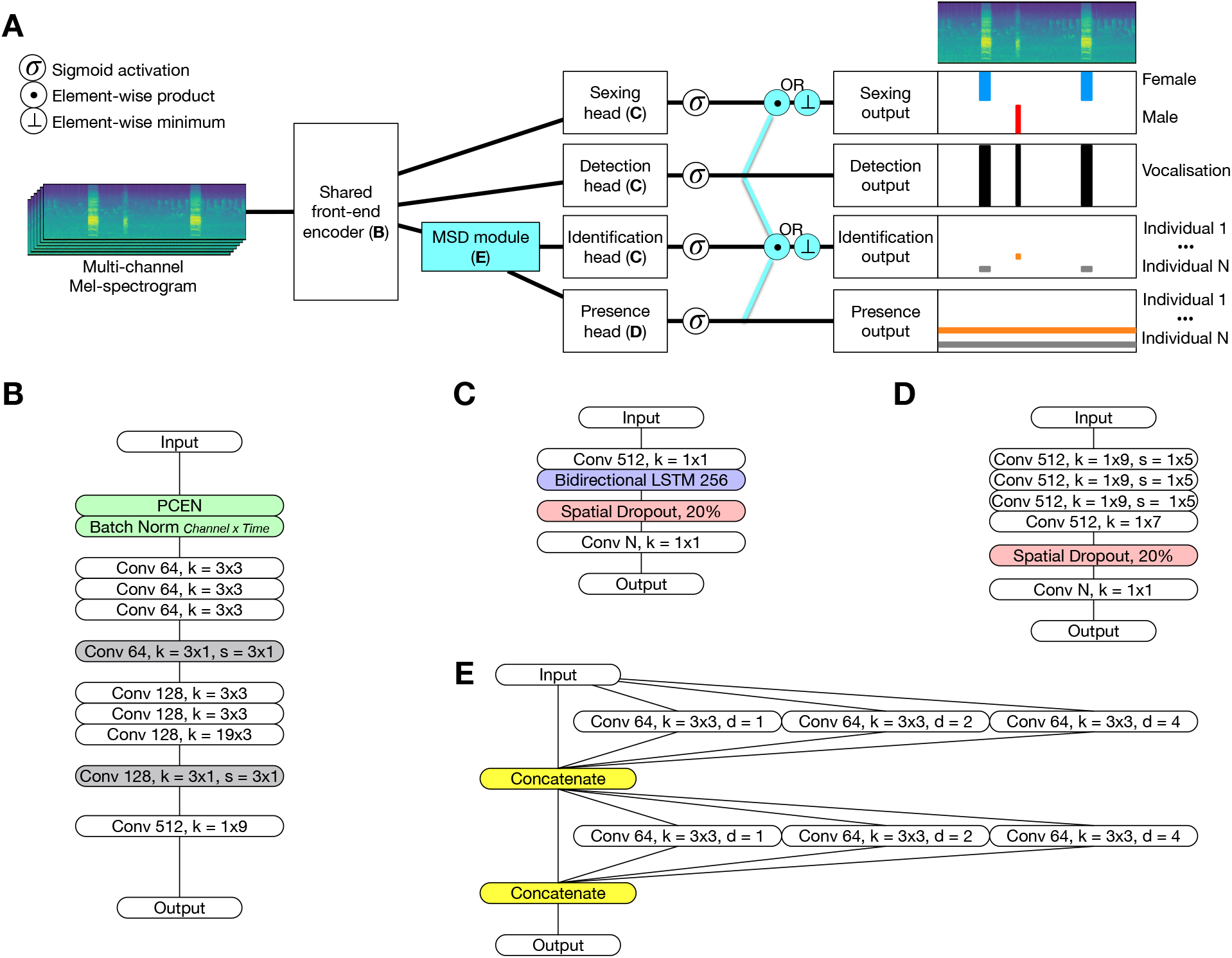
The Rookognise system. A) Overview of the system. Elements in light blue (the MSD module and the two fusion operation, element-wise product and element-wise minimum) are optionally included at training time. On the right: example clip containing three vocalisations, two from a female and one from a male. Under the Mel-spectrogram are the corresponding ground truth labels (in the case of the presence output, stretched in time to match the shape of the other outputs). B) Front-end encoder detail. C) Head detail for the detection, identification and sexing tasks. D) Head detail for the presence task. E) MSD module detail. B-E: boxes represent one layer, with activation layers and Batch Renormalisation layers omitted for readability. Convolution layers (Conv) used the parameters in each box in the following manner, taking for instance the first grey Conv box in B: Conv 64, k=3×1, s=3×1 means a convolution with 64 filters of size 3×1 (3 along the frequency axis, 1 along the time axis) and stride 3×1 (likewise). Boxes coloured according to operation type (green: normalisation, white: convolution, grey: pooling, blue: LSTM, red: dropout, yellow: concatenation). N: number of classes for a particular task (detection: 1, sexing: 2, presence and identification: 15).

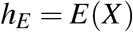

where *E* is the front-end neural encoder, *X* is the input Mel-spectrogram, of dimensions (80, 800, 6), and *h*_*E*_ is the corresponding latent encoding of *X* by *E*, of dimensions (1, 800, 512). *E* has an overall receptive field size of (80, 39, 6), corresponding to the entire frequency axis, slightly under 0.5 s of time, and all audio channels.

#### Multi-Scale Densely-connected (MSD) module

Salient information for individual identification may be present at multiple scales from local to vocalisation-wide. Several mechanisms exist to integrate multi-scale information, but we chose a module proposed by Pelt and Sethian (2017) for its simple design, which the authors note make it applicable to a wide variety of tasks aside from their original task in medical imaging. This module is built by stacking *M* successive blocks, each with *L* parallel dilated convolutional layers with identical parameters except for the dilation rate *d*. We build the module with *M* = 2 blocks and *L* = 3 convolutions per block, using dilation rates *d* of 1, 2, and 4 (Fig. 2E), although we did not extensively experiment with these settings. All convolutions use the same parameters except for this dilation rate: 64 filters, 1×3 filter size. The module is placed directly after the front-end. Formally, this corresponds to the equation:

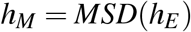

where *MSD* is the module and *h*_*M*_ is the latent encoding of *h*_*E*_ by *MSD*, of dimensions (1, 800, 512).

#### Task-specific heads

We implemented the multi-task learning objective by defining heads with dedicated architecture for each classification task. All heads received the output of either the front-end or the MSD module. The detection, sexing and identification head shared the same architecture (Fig. 2C) including a pointwise convolution layer, a spatial dropout layer, a bidirectional LSTM layer, and a final pointwise convolution. All three heads thus preserved time resolution. The presence head summarised over the time axis using three convolutions with kernel size 1×9 and stride 1×5, followed by a convolution summarising along the remaining time axis information, and finally a pointwise convolution. All heads end with a sigmoid activation for the specific classification task. Formally, this corresponds to:

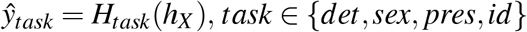

such that *ŷ*_*det*_ is the output of the detection head *H*_*det*_ of dimensions (1, 800, 1), *ŷ*_*sex*_ is the output of the sexing head *H*_*sex*_ of dimensions (1, 800, 2), *ŷ*_*pres*_ is the output of the presence head *H*_*pres*_ of dimensions (1, 1, 15) (15 corresponding to the number of individuals in the study group), and *ŷ*_*id*_ is the output of the identification head *H*_*id*_ of dimensions (1, 800, 15). *h*_*X*_ can be the encoded output of either the front-end *h*_*E*_ or the MSD module *h*_*M*_.

#### Fusing decisions

We finally investigated whether explicitly combining the outputs of different heads could facilitate learning. For instance, combining detection and identification may allow the latter to only attend to frames containing vocalisations, as frames with low detection outputs are unlikely to contain vocalisations. This combination operation was always added after the sigmoid at the end of each head. We tested two operations: element-wise minimum and element-wise product. In our experiments, we only fused outputs with shapes naturally compatible with element-wise operations: 1) detection, presence and identification, and 2) detection and sexing (Fig. 2A).

### 2.4 Implementation details

In this section we detail the implementation of the neural network. We first describe non-standard elements used in the architecture, followed by the regularisation strategies used to prevent overfitting, and finally the loss function and the optimizer algorithm used to actually train the network.

#### Non-standard architecture elements

The field of deep learning has been extremely active in recent years, with both new applications and new architecture elements regularly proposed to improve previous results. We selected some of these elements for their theoretical benefits on performance, briefly described below, and included them in the final system after experimentally validating these benefits. As these elements are not part of our contributions, we do not focus on them further; however, in Table A.2 we show how using these non-standard elements improved the system compared to their standard counterparts. First, we used the Mish activation function instead of the ReLU family of activations, for its continuous derivability and self-regularising property, that prevent gradient vanishing and thereby improve training (Misra, 2019). Second, we used Batch Renormalisation (Ioffe, 2017) instead of Batch Normalisation. Batch Normalisation improves training efficiency and adds robustness to specific initialisation quirks (Ioffe and Szegedy, 2015), but suffers when batches are not independent and identically distributed, which Batch Renormalisation aims to correct. Third, we used strided convolutional pooling (Pankajakshan et al., 2018) instead of max pooling. Max pooling essentially discards the majority of the data when downsampling since only the highest value for each pooling operation is kept. We expected this may learning, as individual identity may be encoded by fine-grained cues. Therefore, we chose strided convolutional pooling to avoid discarding any information during training.

#### Regularisation

Regularisation prevents the network from overfitting the training data, which risks losing generalisation ability to unseen data. We used several of these techniques in our experiments. First, we used early stopping with a waiting period of 15 epochs, to directly stop training when validation data performance no longer improved. The waiting period was based on the learning rate cycle (see below), to allow at least two minima of the learning rate to occur before stopping. Second, we used spatial dropout with a rate of 20% before the final layer of each head. Spatial dropout prevents overfitting by randomly dropping entire feature maps at each iteration of training, which prevents feature co-adaptation that could lead to overfitting (Srivastava et al., 2014). Third, we applied L2-norm weight decay with a strength of 10^*−*4^. This penalises large absolute values of the weights that can destabilise training (Smith, 2018). Fourth, we used label smoothing so that the actual training labels used to minimise the loss functions were not 0 and 1 but 0.05 and 0.95, respectively. Label smoothing similarly encourages lower weights for stable training, particularly in the last layer of the network (Szegedy et al., 2016).

#### Objective function and optimizer

All networks were trained to minimise a focal binary cross-entropy loss (Lin et al., 2020, FL,) with the formula:

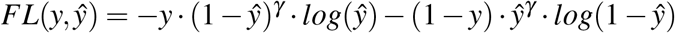

where *y* is a ground truth label, *ŷ* is the corresponding network prediction, and *γ* is a focusing parameter, which reduces the contribution of correctly classified samples to the loss. When *γ* = 0, FL reduces to the standard binary cross-entropy. We chose *γ* = 2 in our experiments as it improved over the standard binary cross-entropy (see Table A.2 for details). The mean of the FL over all samples was used for each head. In the multi-task configurations, we used the unweighted sum of the loss corresponding to all four tasks:

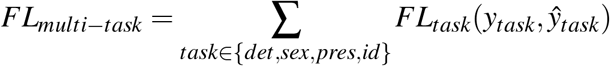

We used the Ranger21 optimiser (Wright and Demeure, 2021), with weight decay 10^*−*4^. Ranger21 serves to solve some issues with more usual optimisers such as Adam, which can be unstable at the beginning of the training process and so derail the training procedure, along with integrating a number of heuristics which improve various aspects of training (see Table A.2 for a comparison between Ranger21 and Adam). We chose a cyclical learning rate schedule (Smith, 2018), linearly interpolating between 10^*−*6^ and 10^*−*3^ with cycles of 6 epochs, and halved the maximum learning rate after each cycle. Cyclical learning rates can greatly accelerate convergence by allowing the network to update much faster when the learning rate grows, and converge to more precise minima of the loss when the learning rate decreases. Finally, we used mixed precision training (Narang et al., 2018) to accelerate training and decrease memory footprint.

#### 2.5 Training procedure

The networks were trained for a maximum of 50 epochs to minimise the FL on the Mel-spectrograms of the multichannel recordings. We semi-randomly distributed the annotated sessions at the start of the experiments. We adjusted the distribution in the training data so all individuals were represented as equally as possible, although this also means not all individuals could be represented in the validation and test data (Fig. 1). The annotation files were not split during either training or evaluation, to avoid data leakage due to the way vocalisations were sampled during training.

For the training data, clips were randomly sampled around each vocalisation so as to include this vocalisation in its entirety (although other vocalisations in the same clip could be cut off). This meant the network was unlikely to ever receive the same spectrogram twice, encouraging it to learn the underlying structure of the data. Additional clips were randomly sampled from background-only parts of the recordings to increase the variety of background noises. For the validation and the test data, the recordings were simply split into adjacent, non-overlapping clips and sent in order to the network.

Vocalisations from the Unknown or Multiple classes were excluded from training by assigning them a weight of 0 during loss computation. The detection task was exempt from this, as identity is irrelevant to whether or not a vocalisation occurs.

Finally, we used the same random seed to initialise all runs for fair comparison. This meant all networks started from the same initial weights, and the training data was always fed to the network in the same order at a given epoch, so all differences between networks should be due to differences in architecture.

#### 2.6 Experiments

We investigated and compared several configurations to build the final network, following the description in section 2.3. For clarity, all configurations were named using the nomenclature in Table 1. First, single-task networks were trained and evaluated to form a baseline. For each sub-task, one network was trained without and with the MSD module. Then, we trained the multi-task network, using the best configurations of each sub-task from the single-task experiment. Following this, we included either of the two fusion operations described above in the multi-task network. As a final test, we investigated the impact of using multi-channel audio compared to using only one channel. We hypothesised that multi-channel audio is more efficient due to including additional information that the network may access, such as delays between channels or simple differences in distance: while with all six microphones the maximum distance between bird and microphone was only about 10 m, with one channel distance to microphone it might increase up to about 26 m. The configuration corresponding to this case of using a single audio channel was named the Rookognise RC (Random Channel) configuration.

**Table 1:** Description and nomenclature of the configurations in the experiments. From left to right: Configuration is the name used in the text for each network, the Task learning column denotes whether the network learned a single task or used the multi-task approach, the Task column corresponds to the task(s) learned in a given configuration, the MSD module column corresponds to whether the MSD module is included or not in a given configuration, and the Combination column refers to the element-wise operation used for combining decisions in the configuration, if any.

| Configuration | Task learning | Task | MSD module | Combination |
| --- | --- | --- | --- | --- |
| ST_Det | Single | Detection | No |  |
| ST_Det_MSD | Single | Detection | Yes |  |
| ST_Sex | Single | Sexing | No |  |
| ST_Sex_MSD | Single | Sexing | Yes |  |
| ST_Pres | Single | Presence | No |  |
| ST_Pres_MSD | Single | Presence | Yes |  |
| ST_Id | Single | Identification | No |  |
| ST_Id_MSD | Single | Identification | Yes |  |
| MT_None | Multi | All | Yes |  |
| MT_Product | Multi | All | Yes | Product |
| MT_Min | Multi | All | Yes | Minimum |

#### 2.7 Evaluation

We ranked our models using the Area Under the Receiver Operating Curve (AUROC) and the Area Under the Precision-Recall Curve (AUPRC). AUROC and AUPRC both summarise the performance of a network over its entire output range. AUROC indicates the probability a classifier will score a random positive sample higher than a random negative sample. No matter the dataset, random guessing leads to 50% AUROC and perfect classification to 100% AUROC (Fawcett, 2006). However, AUROC can over-emphasise the correct rejection of negative samples, especially if negative samples outnumber positive samples, in which case AUPRC can be more representative (Saito and Rehmsmeier, 2015). AUPRC is based on precision (the ratio of true positives over predicted positives) and recall (the ratio of true positives over labelled positives), and evaluates the ability of the classifier to correctly classify positive samples without false positives or negatives, and ignores true negatives (i.e. correct rejections). Unlike AU-ROC, AUPRC is not independent of the proportion of positive samples in the dataset, and so cannot be compared between different datasets. During our experiments, we saved the models at the epoch with the highest AUPRC, as computed on the validation dataset. In the multi-task configurations, only the identification task validation AUPRC was used. The reported values of AUROC and AUPRC were the average of the AUROC and AUPRC associated with each class. The average was computed from the classes present in a given dataset (i.e., only 13 of the 15 birds were present in the test dataset, so the corresponding results were averaged over 13 classes instead of 15).

The ROC and PRC curves represent the performance of the network at various decision thresholds. The AUROC and AUPRC metrics integrate over all possible threshold values; as such, while they are convenient to evaluate network performance, they are not sufficient when we are interested in selecting a threshold when we apply the network to actual unseen data. To do so, we used the curves themselves. Since our dataset is heavily imbalanced, we used the AUPRC curve, and we chose to directly plot precision and recall as a function of the threshold value for clearer visualisation.

## 3 Results

We first built the Rookognise system by comparing single-task networks with and without the MSD module, before combining the best configurations into a single multi-task network. We then compared the multi-task learning-only approach to the approaches incorporating output fusions, to arrive at the final system. We briefly investigated the benefit of using the multi-channel audio dataset compared to a mono-channel version of the dataset. Finally, we illustrated the use of the Rookognise system on unseen data from the test dataset.

### 3.1 Integrating multi-scale information with the MSD module

In this section we investigated the benefits of multi-scale information integrated by the MSD module. We established the baseline performance of the system by training single-task networks dedicated to each of the four sub-tasks, with and without the MSD module (Table 2). In the following, we report results on the test data: positive numbers where the MSD module improved performance, and negative numbers otherwise. For the detection sub-task, ST Det outperformed ST Det MSD (−7.34% AUROC, −3.23% AUPRC). For the sexing sub-task, ST Sex outperformed ST Sex MSD (−0.07% AUROC, −0.01% AUPRC). For the presence sub-task, ST Pres MSD outperformed ST Pres (+2.88% AUROC, +1.28% AUPRC). Finally, for the identification sub-task, ST Id MSD outperformed ST Id (+0.94% AUROC, +3.22% AUPRC). Overall, the MSD module improved performance for the tasks involving more complex classification (in this case, with higher number of possible outputs). Though the improvement was not systematic for all the sub-tasks, the integration of multi-scale information in the MSD module likely improved the ability of the network to disentangle important information at both local and global scales to identify specific individuals. Overall, the MSD module efficiently captured multi-scale time-frequency information in audio signals for complex tasks. On the other hand, the front-end encoder alone was sufficient to detect rook vocalisations against all other sounds, and to classify rook vocalisations by sex, likely simpler tasks to learn.

**Table 2:** Test data performance of the networks, at the best epoch determined by validation data AUPRC. For each column, the best performance is emphasised in bold (separately for single-task and multi-task configurations). Underlined values represent where the multi-task configurations outperformed the corresponding best single-task network. The last line corresponds to the mono-channel configuration, obtained by training the network using only one of the microphones randomised for each audio clip.

| Configuration | Detection |  | Sexing |  | Presence |  | Identification |  |
| --- | --- | --- | --- | --- | --- | --- | --- | --- |
|  | AUROC | AUPRC | AUROC | AUPRC | AUROC | AUPRC | AUROC | AUPRC |
| ST_Det | **94.03** | **81.04** |  |  |  |  |  |  |
| ST_Det_MSD | 86.69 | 77.81 |  |  |  |  |  |  |
| ST_Sex |  |  | **98.88** | **89.72** |  |  |  |  |
| ST_Sex_MSD |  |  | 98.81 | 89.71 |  |  |  |  |
| ST_Pres |  |  |  |  | 83.62 | 53.11 |  |  |
| ST_Pres_MSD |  |  |  |  | **86.50** | **54.39** |  |  |
| ST_Id |  |  |  |  |  |  | 92.87 | 55.28 |
| ST_Id_MSD |  |  |  |  |  |  | **93.81** | **58.50** |
| MT_None | 89.46 | 76.35 | 97.83 | 87.26 | <u>87.97</u> | <u>62.73</u> | <u>94.50</u> | <u>60.86</u> |
| MT_Min | <u>94.69</u> | <u>81.72</u> | 97.64 | 87.47 | **89.94** | **64.65** | 93.47 | <u>61.30</u> |
| MT_Product (Rookognise) | **96.95** | **89.31** | **98.21** | **88.95** | <u>88.03</u> | <u>63.64</u> | **96.11** | **63.06** |
| Rookognise_RC | 92.35 | 81.78 | 95.04 | 81.65 | 84.15 | 58.70 | 93.70 | 59.29 |

### 3.2 Multi-task learning and fusing sub-task decisions

In this section, we investigated the impact of multi-task learning. We compare the best single-task networks determined in section 3.1 with the multi-task configurations MT None, MT Product, and MT Minimum (Table 2, middle part). In this case, we report positive numbers where the MT configurations outperformed the ST networks, and negative numbers otherwise. The MT None configuration already outperformed the single-task configurations on the presence (+1.47% AUROC, +8.34% AUPRC) and identification (+0.69% AUROC, +2.36% AUPRC) sub-tasks) sub-tasks, though not on the detection sub-task (−4.57% AUROC, −4.69% AURPC). Adding either fusion operation maintained or even amplified the improved performance on presence and identification, and also improved detection performance. In particular, MT_Product obtained greatly improved performance on all three sub-tasks compared to any other configuration. Interestingly, none of the MT configurations outperformed the single-task networks on the sexing sub-task, although MT Product was only slightly worse (−0.67% AUROC, −0.77% AUPRC). Overall, the multi-task approach greatly improved learning ability on the more complex tasks. Fusing the decisions of multiple sub-tasks further boosted performance, especially in the detection sub-task which did not benefit from the multi-task approach alone. In particular, using the element-wise product resulted in the strongest overall performance. Following these experiments, we selected the MT Product configuration as the final system, which hereafter corresponds to the ”Rookognise” system.

### 3.3 The impact of multi-channel audio

Using multi-channel audio includes a lot more information than using only one audio channel. As just one example, vocalisations clearly audible on one channel may not be audible on another (Fig. 3). In a final experiment, we investigated to what extent the Rookognise system may benefit from the multi-channel aspect of the training data. To do so, we simply re-trained the system in the Rookognise configuration, using a single-channel version of the dataset obtained by randomly sampling one audio channel for each sample, resulting in the Rookognise RC configuration (Table 2, last line). This configuration obtained lower performance than the Rookognise system on all sub-tasks (detection: −4.59% AUROC, −7.53% AUPRC; sexing: −2.94% AUROC, −7.30% AUPRC; presence: −3.89% AUROC, −4.94% AUPRC; identification: −2.41% AUROC, −3.7% AUPRC). This shows the ability of the network to efficiently capture additional (e.g. spatial) information as provided by the multi-channel microphone array, in particular for the detection of rook vocalisations. Nevertheless, Rookognise RC still obtained reasonable performance despite using a single audio channel, making it usable if storage space for the audio recordings is a concern.

**Figure 3:**
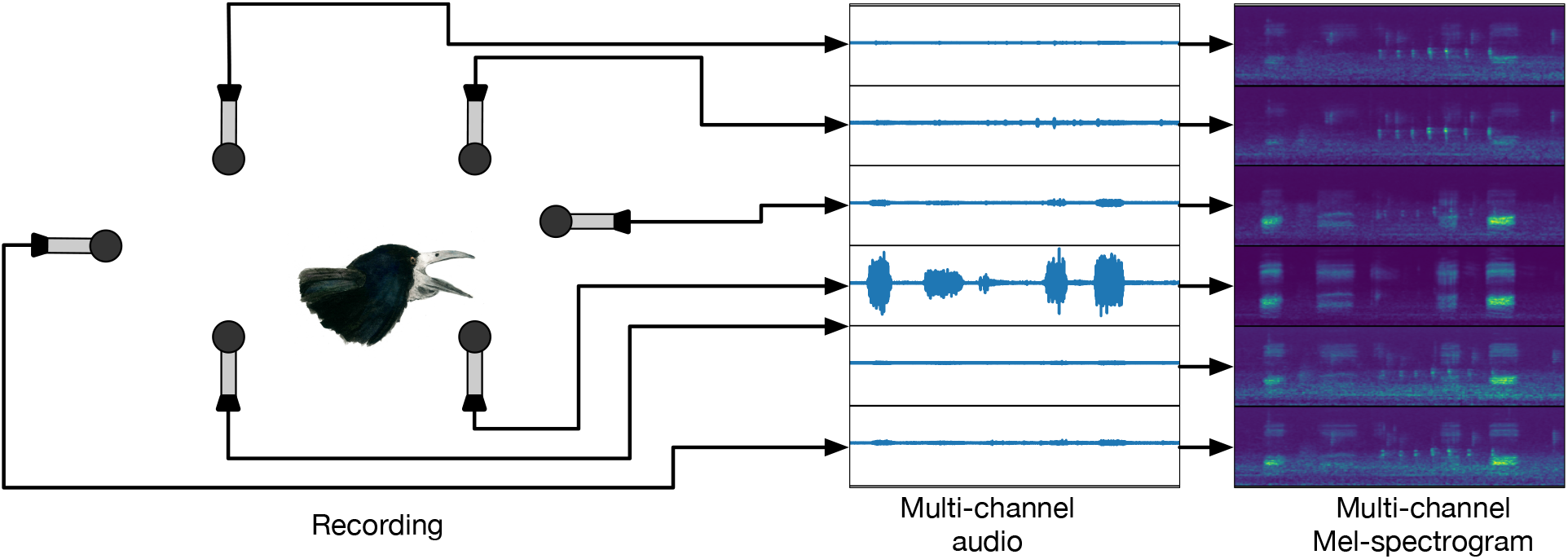
Schematic illustration of the multi-channel setup. Here, a rook is vocalising closest to the rightmost microphone (left), corresponding to the fourth channel from the top in the audio recording (middle) and in the corresponding Mel-spectrograms (right). On the other hand, the leftmost microphone, furthest from the rook, barely records anything over the background noise

### 3.4 Detecting vocalisations with the Rookognise system

We examined in further detail the performance of the Rookognise system on unseen data, starting with the detection of rook vocalisations from raw recordings. We computed the precision (ratio of true positives to predicted positives) and recall (ratio of true positives to labelled positives) metrics, as well as the proportion of the recording data retained as a function of threshold value (between 0 and 1, corresponding to the output range of the system) for each sub-task (Fig. 4). For the detection sub-task (Fig. 4, upper left), a low threshold of 0.05 was already enough to discard approximately 75% of the data, i.e. most background noise could be easily eliminated. This low threshold missed very few vocalisations (95.0% recall) but also falsely classified some noise as rook vocalisations (35.9% precision). Conversely, a higher threshold of 0.49, computed to optimise the F1-score (the harmonic mean of precision and recall), had much better precision (90.1%) and retained only 8.5% of the data, at the cost of lower recall (79.2%). A good threshold value would of course depend on the user, but optimal thresholds may lie between those two values to balance human effort reduction (with a high threshold) and false detections (with a low threshold).

**Figure 4:**
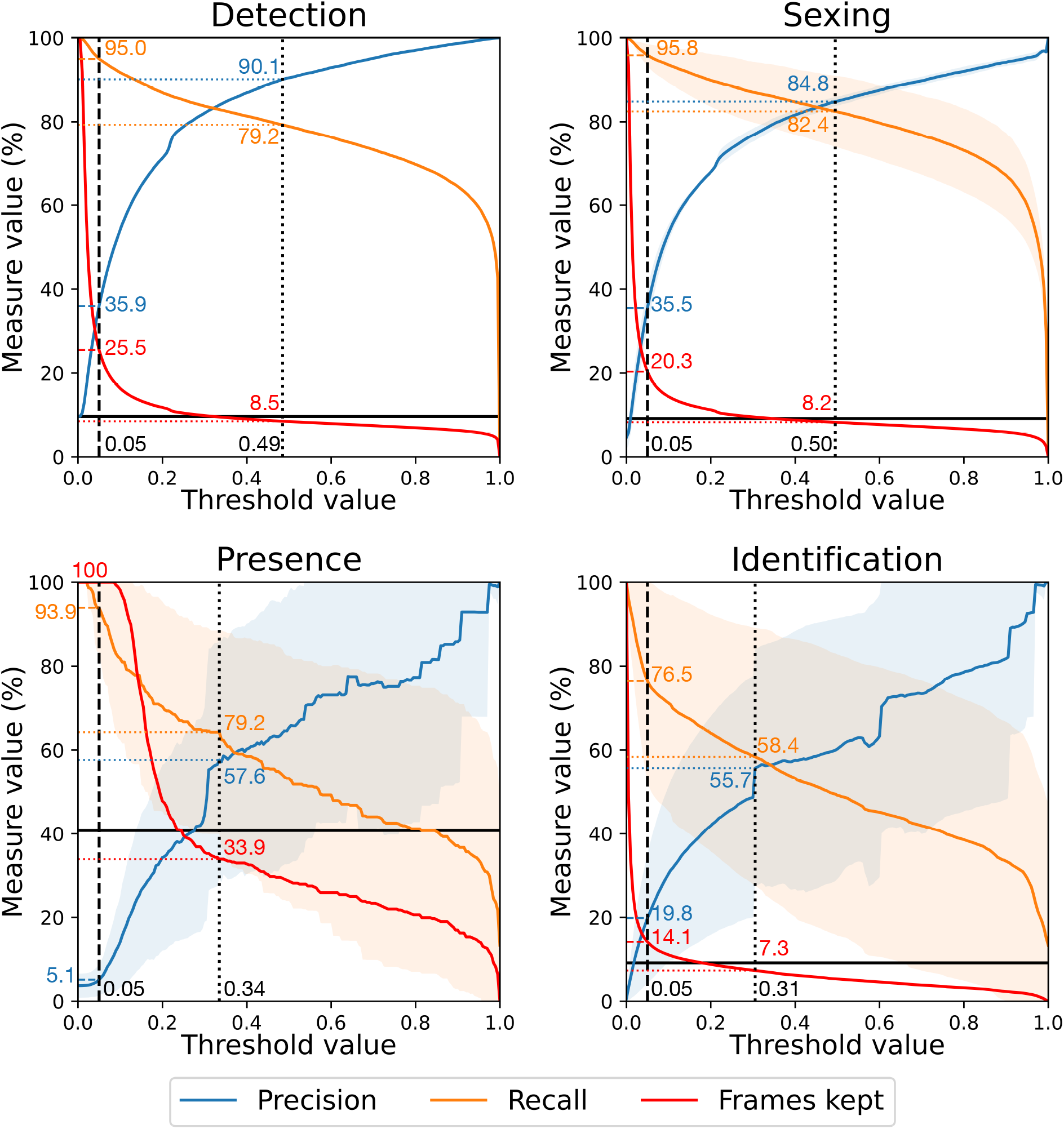
Test performance of the Rookognise system for each sub-task. Performance is measured using precision, recall and proportion of spectrogram frames kept (as a proxy of human effort reduction). For the sexing, presence, and identification sub-tasks the precision and recall curves are averaged over classes and the shaded area represents one standard deviation away from this average. In all plots, the black vertical lines corresponds to specific threshold values: dashed lines correspond to the conservative threshold at 0.05, dotted lines to the threshold optimising F1-score (the harmonic mean of recall and precision). Coloured horizontal lines are the values of all three metrics at the corresponding thresholds, with the associated values in the corresponding colours.

### 3.5 Identifying emitters with the Rookognise system

We finally directly evaluated how well the network identified individual rooks. For the sake of this evaluation, we used only predictions corresponding to vocalisations from identified individuals in the test labels (i.e. we discarded all Unknown or Multiple class vocalisations, as well as overlapping vocalisations). Each vocalisation frame was then attributed to the individual with the highest predicted score. We computed the confusion matrix from the resulting predictions, as well as the corresponding precision and recall associated with each individual (Table 3). The Rookognise system had strong overall performance identifying individuals, attributing 70.6% of vocalisation frames to the correct individual, with an average precision of 65.2% and average recall of 64.7%. Seven individuals in particular were identified with both precision and recall over 75%. Several individuals (O, P, and E) were more poorly identified. While these individuals were among the less represented in the data (e.g. O has 428 vocalisations in the training dataset), other individuals (e.g. S with 239 vocalisations and J with 125 vocalisations) were much better identified (S: 89.5% precision and 78.5% recall; J: 84.2% precision and 76.8% recall). In general, we found no correlation between identification performance (whether measured as precision or recall) and number of vocalisations in either the training or test data (four Pearson correlation tests: either precision or recall, correlated with number of vocalisations in either training or test data, all *p >* 0.05). We further examined two noteworthy cases.

**Table 3:** Confusion matrix computed from the test data predictions. The predictions were restricted to labelled vocalisations, with overlapping and non-attributed vocalisations removed. Birds are coloured by sex (males in blue, females in red). Columns correspond to labels, rows to predictions. For instance: for the cell in the first row (A), second column (B) the network predicted 2 frames as produced by A that were actually produced by B. Diagonal cells (in grey) correspond to correct identifications. For each bird, the second-to-last column indicates predicted frames, the last column indicates precision, the second-to-last row indicates labelled frames, and the last row indicates recall. The four bottom-right cells contain: total frames (top left), average recall (bottom left), average precision (top right) and overall accuracy (bottom right). Individuals that were absent from the test dataset (F and R) had their recall and precision left blank (note that recall is undefined and precision is 0 in the absence of positive samples) and did not contribute to the averages.

|  | A | B | E | F | G | H | J | K | L | M | O | P | R | S | T | Predicted | Precision |
| --- | --- | --- | --- | --- | --- | --- | --- | --- | --- | --- | --- | --- | --- | --- | --- | --- | --- |
| A | 1009 | 2 | 0 | 0 | 0 | 8 | 0 | 130 | 0 | 71 | 0 | 1 | 0 | 0 | 0 | 1221 | 82.6 |
| B | 14 | 3943 | 111 | 0 | 0 | 123 | 0 | 8 | 0 | 12 | 92 | 0 | 0 | 0 | 918 | 5221 | 75.5 |
| E | 0 | 8 | 437 | 0 | 0 | 0 | 0 | 81 | 4 | 204 | 0 | 0 | 0 | 0 | 124 | 858 | 50.9 |
| F | 0 | 0 | 13 | 0 | 18 | 0 | 0 | 0 | 53 | 0 | 0 | 0 | 0 | 0 | 0 | 84 | - |
| G | 1 | 22 | 41 | 0 | 2729 | 5 | 0 | 748 | 45 | 0 | 0 | 26 | 0 | 27 | 0 | 3644 | 74.9 |
| H | 4 | 0 | 106 | 0 | 0 | 3957 | 0 | 2078 | 6 | 0 | 0 | 0 | 0 | 0 | 20 | 6171 | 64.1 |
| J | 0 | 0 | 0 | 0 | 0 | 0 | 251 | 0 | 45 | 0 | 0 | 0 | 0 | 2 | 0 | 298 | 84.2 |
| K | 3 | 13 | 159 | 0 | 0 | 35 | 0 | 5541 | 3 | 8 | 5 | 0 | 0 | 0 | 934 | 6701 | 82.7 |
| L | 0 | 9 | 210 | 0 | 11 | 0 | 76 | 4 | 1583 | 0 | 0 | 196 | 0 | 15 | 0 | 2104 | 75.2 |
| M | 0 | 3 | 0 | 0 | 9 | 0 | 0 | 213 | 0 | 1783 | 372 | 0 | 0 | 0 | 65 | 2445 | 72.9 |
| O | 0 | 0 | 64 | 0 | 0 | 0 | 0 | 676 | 1 | 73 | 33 | 0 | 0 | 0 | 0 | 847 | 3.9 |
| P | 0 | 0 | 73 | 0 | 0 | 0 | 0 | 31 | 0 | 0 | 0 | 81 | 0 | 0 | 0 | 185 | 43.8 |
| R | 0 | 0 | 0 | 0 | 0 | 0 | 0 | 0 | 21 | 0 | 0 | 16 | 0 | 0 | 0 | 37 | - |
| S | 0 | 0 | 0 | 0 | 21 | 0 | 0 | 0 | 0 | 0 | 0 | 0 | 0 | 179 | 0 | 200 | 89.5 |
| T | 0 | 7 | 214 | 0 | 0 | 402 | 0 | 24 | 0 | 0 | 0 | 0 | 0 | 4 | 439 | 1090 | 40.3 |
| Labelled | 1031 | 4007 | 1428 | 0 | 2788 | 4530 | 327 | 9534 | 1761 | 2151 | 502 | 320 | 0 | 227 | 2500 | 31106 | 64.7 |
| Recall | 97.9 | 98.4 | 30.6 | - | 97.9 | 87.4 | 76.8 | 58.1 | 89.9 | 82.9 | 6.6 | 25.3 | - | 78.9 | 17.6 | 65.2 | 70.6 |

The first case was the misattribution of some of K’s vocalisations to H (Table 3, 6th row, 8th column), which was almost exclusively due to the misidentification of one long song-like bout produced by K. These two males, especially H, were also the ones who produce the longest song-like bouts.

The second case was the misattribution of almost all of O’s vocalisations to M (Table 3, 9th row, 10th column), resulting in O being by far the worst-identified individual in the group at only 3.9% precision and 6.6% recall. These two males shared several vocalisation types produced by not other individual, and to our ear, they sounded very similar in general. However, this was also the case for a pair of females, J and L, but these females were successfully identified by the network (J: 84.2% precision, 76.8% recall; J: 75.2% precision, 89.9% recall).

### 3.6 Post-processing decisions: an example on the test data

In this final section we combined the decisions determined in the previous sections, along with some simple post-processing steps which can improve the accuracy of the Rookognise predictions. We then separately compared their effects on the test data labels. First, we used the detection threshold of 0.05. Second, following Cohen et al. (2022), we discard short predictions that are likely false positives; we set the threshold at 5 successive frames, corresponding to 0.0625 s at our time resolution (Fig. 5A). The 5 frames threshold effectively discarded 37% of false positive detections, at the cost of removing 1.5% of the labelled vocalisations. This had relatively minor effect on the detection precision (37.8% instead of 35.9% without discarding short detections) and recall (94.7% instead of 94.9% without discarding short detections), since we remove relatively few frames in total. Third, we discarded predictions for the identification if the highest predicted score is not at least twice as high as the second highest score (Fig. 5B). This effectively discarded predictions where the network was not sufficiently confident in attributing the vocalisation to an individual, and increased accuracy to 84.7% (against 70.6% without this step) while retaining 72.0% of vocalisation frames. We show two examples of system outputs after applying these steps in Fig. 6. In Fig. 6A, four calls from the female L and one call from the male A were correctly detected, sexed, and attributed to the correct individual. In Fig. 6B, we illustrated a clip containing a long song-like bout produced by H. Again, the network correctly detected, sexed and attributed these vocalisations, including separating most vocalisations even if they occur in quick succession.

**Figure 5:**
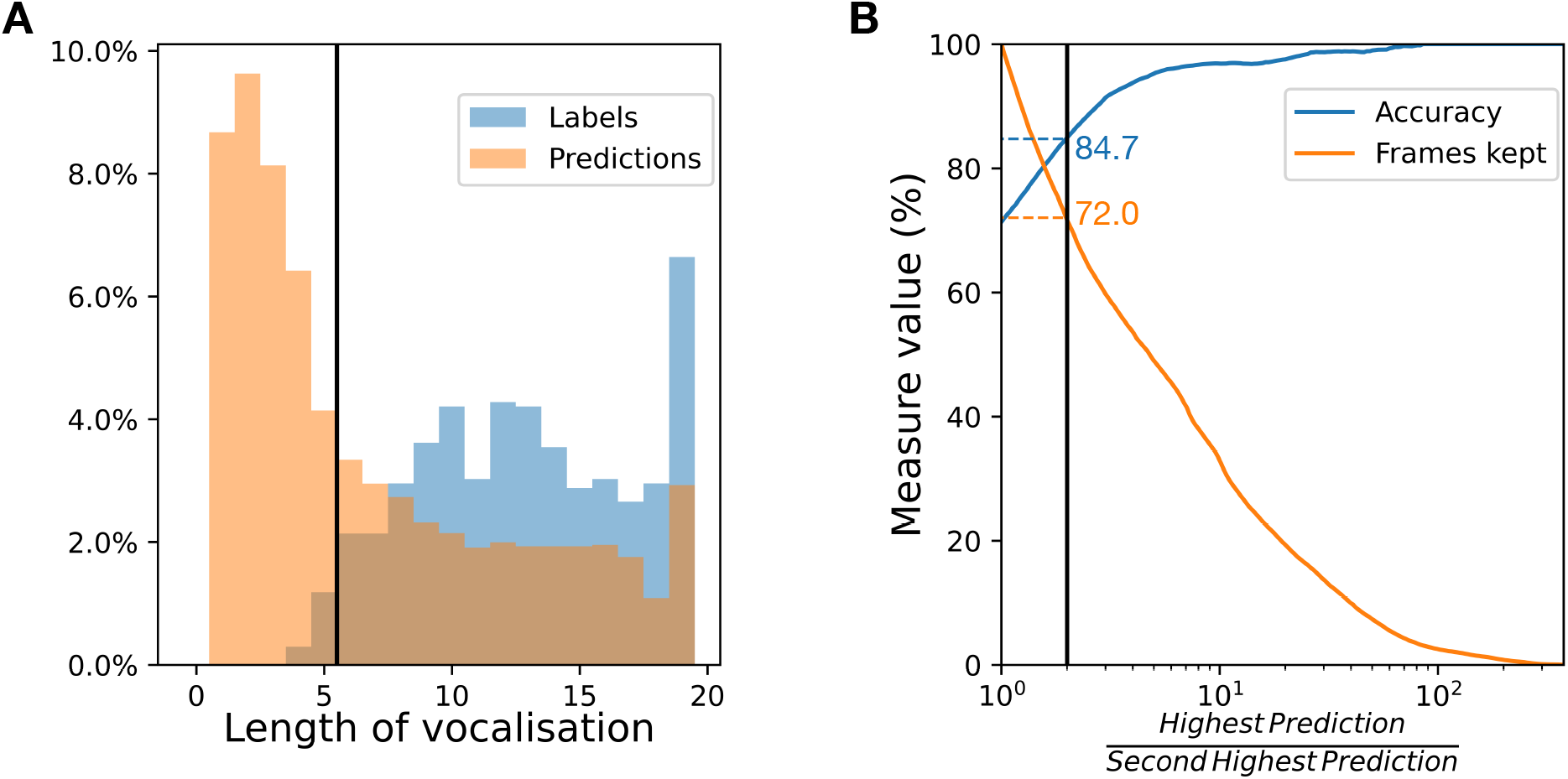
Post-processing of the system predictions. A) Distribution of the Rookognise detection sub-task predictions (orange bars) and of the labelled vocalisations (blue bars) by length in frames. The black line represents the threshold, chosen at 5 consecutive frames. Given the distribution of lengths in the labels, Rookognise predictions shorter than 5 frames are likely false positives and can be safely removed. Only detections (for both labelled and predicted vocalisations) up to 20 frames long are included to avoid crowding the plot. B) Accuracy and proportion of frames retained as a function of the ratio between the highest and second-highest prediction for the identification task. Both measures are computed based on the frames corresponding to labelled vocalisations, as in Table 3. The black line represents the point where the ratio of highest prediction to second-highest prediction is 2. Identification predictions are only retained if the highest prediction sufficiently higher, meaning the network is confident in identifying one individual.

**Figure 6:**
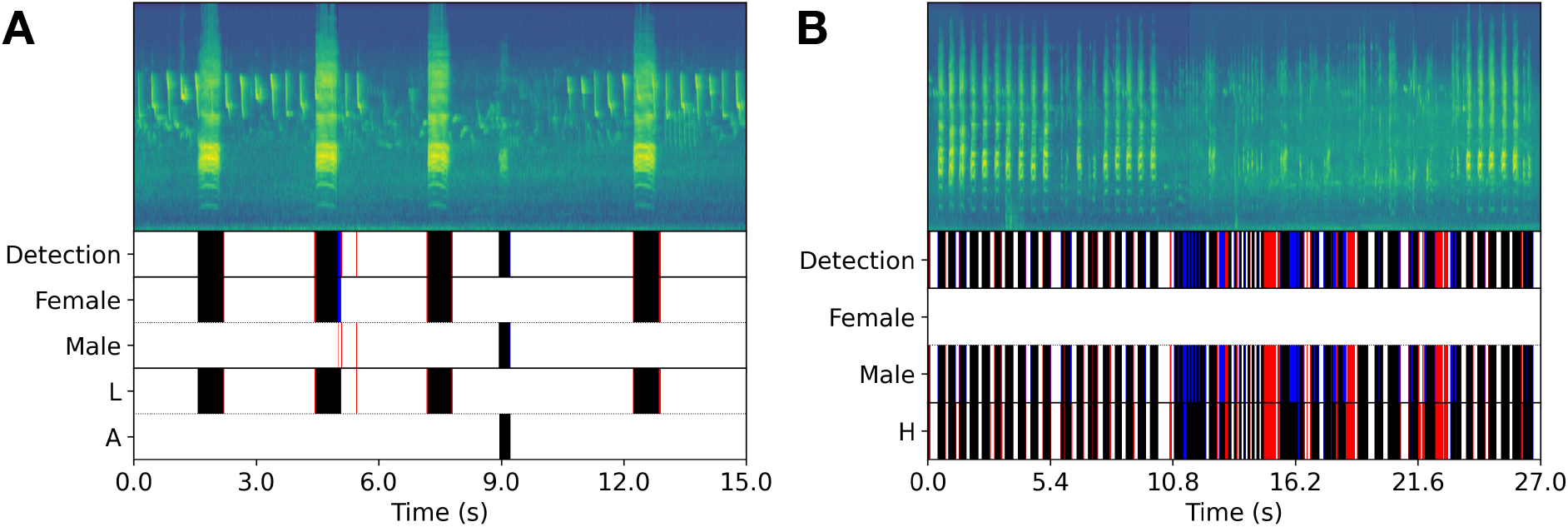
Two example Mel-spectrograms from the test dataset (only one audio channel shown), with corresponding network outputs for detection, sexing, and identification, shown after the post-processing detailed in the text. A) Calls produced by one female (L) and one male (A). B) Song-like bout produced by one male (H). True negatives are in white, true positives in black, false negatives in blue, and false positives in red.

## 4 Discussion

We proposed Rookognise, a deep learning system for the acoustic detection and vocalization-independent identification of rooks in an outdoor soundscape. To our knowledge, the system is the first approach that can acoustically identify individuals regardless of the vocalisation produced. We approach the task by combining three different mechanisms: the multi-scale densely-connected MSD module, the multi-task approach where we divide the complex overall task into a cascade of increasingly complex tasks, and the explicit combination of outputs during training. All three mechanisms had substantial benefits to generalisation performance on unseen data. Multi-task learning has been proposed as acting as a regularisation mechanism (Liebel and Körner, 2018), although naive multi-task approaches may impede learning due to conflicting objectives. This may have caused performance loss on the detection and sexing sub-tasks for the *MT*_*N*_*one* configuration. In addition to the regularising effect of multi-task learning, the auxiliary tasks allow users to separately determine task-specific thresholds for more direct control over the precision and recall trade-off. Furthermore, conditioning the outputs of more complex tasks on the outputs of simple tasks may help the learning process, possibly by acting as an attention or a hierarchical classification (Nolasco and Stowell, 2022) mechanism: in this case the detection and presence tasks allow early rejection of, respectively, non-vocalisations or individuals absent from the audio clip.

The final system had strong overall performance according to all metrics used. Unfortunately, there are as yet few comparable approaches to acoustically identify individuals irrespective of vocalisation types (although there have been approaches to identify individuals by image data in multiple species, including birds). Stowell et al. (2019a) reported highly successful individual identification in three bird species, the little owl (*Athene noctua*), the chiffchaff (*Phylloscopus collybita*), and the tree pitpit (*Anthus trivialis*). They used a Random Forest classifier attributing an entire audio file to one individual, with high AUROC (85-94%, slightly lower than Rookognise). However, their metric was computed as one decision per file and their dataset was by design restricted to one focal individual per file, so the comparison with our system is by necessity rather limited. Comparing different systems is generally an issue in deep learning, as many tasks lack a standardised benchmark with a well-defined dataset and associated metrics, to allow different approaches to be compared rigorously. For individual acoustic identification, this issue is even more prevalent due to the current absence of similar studies (Stowell, 2022). Nevertheless, Rookognise appears well suited to monitoring a social group in less controlled conditions, i.e. outdoors with multiple individuals in proximity.

The strong performance of the system was diminished in a few cases. From examining predictions on the test data, these errors consisted mostly of: 1) boundary errors, most often due to the network slightly overestimating the times spanned by vocalisations; 2) some false positives due to the acoustic properties of specific percussive noises, which share similar structure to some very short vocalisations produced by some of the females, and 3) identification errors, mostly confusions between specific individual pairs (e.g. K with H, O with M). In both cases, we could identify the source of the misattribution. In the K/H case, a long vocalisation bout produced by K was attributed to H. The system may have learned to preferentially attribute long song-like sequences to H, because he is the one with longest bouts. Other cases of identification errors may come from vocal similarity between the individuals in the study group. In the O/M case, for example, O and M shared their most common vocalisation, which accounted for a majority of O’s vocalisation in the test data. Note that the system did not otherwise identify individuals proportionally to their dataset presence (i.e. individuals that vocalised less were not less correctly identified).

In many bird species, social proximity and vocal similarity are positively correlated (Brown, 1985; Grießmann and Naguib, 2002; Hausberger et al., 1995), but in our case the misidentified pairs did not correspond to a particularly close social bonds (Boucherie et al., 2016), while individuals involved in other bonds were not confused by the network (Table S.1). Individuals from similar genetic or cultural origins (same colonies) may also exhibit vocal similarities (Lemasson et al., 2011). In our group, two individuals (E and J) were genetically related, but they were not substantially confused by the network (Table A.1). Even if some birds may share vocal similarities due to these factors (social, genetic or cultural proximity), the system still correctly classified identities.

At present, the Rookognise system necessitates knowing the individuals present in the study group to build the network and actively identify each individual. Unknown individuals cannot be identified by the system, but the system is designed to filter these unknown individuals out if desired: vocalisations from unknown individual rooks can be detected by the system (i.e. receive high scores from the detection sub-task) but without being attributed to an individual (i.e. receive low scores from the identification sub-task). As such, the Rookognise system can already be used as is to detect vocalisations from another group of rooks, which can greatly facilitate its deployment to other groups, i.e. only requiring to update the individual identification sub-task while other sub-tasks are already trained. More generally, we believe that the network can be deployed to monitor groups of any other species: birds, primates, cetaceans, provided that the number and identity of individuals in the group are known, and, of course, that the species produces vocalisations as part of its communication. More than a proof of concept, the system should be adaptable to any species no matter how harmonic their vocalisations are. Indeed, the system is already successful on rooks, birds that are characterised by their harsh, inharmonic calls like many corvids (Fletcher, 2000). Neural networks often generalise well, although the risk of overfitting to a particular problem is always present. For instance, Stowell et al. (2019a) pointed out that a classifier trained to recognise one particular individual associated with a particular background noise, may fail to recognise the same individual in a different background noise. Our system was designed to limit this by explicitly training on background noise samples. In addition, the dataset itself, due to the complexity of the background noises recorded in an outdoors setting, likely limits the potential for overfitting on this type of data. Finally, none of the decisions involved in the design of the system were specifically tailored to the species or the group. The main limit to deploying the system to the study of another species is that said species must vocalise frequently enough to collect a sufficient amount of vocalisations in a reasonable time.

Extending the system to conditions where the number of individuals is not known *a priori* requires reformulating the identification task as an open-set recognition problem, a much more technically difficult task. Previous approaches to this problem have largely been in the form of speaker verification (Ntalampiras and Potamitis, 2021; Ptacek et al., 2016; Stowell et al., 2019a) or clustering approaches including deep learning-based ones (Goffinet et al., 2021; Kirschel et al., 2009; Pagliarini et al., 2021) to determine whether vocalisations are produced by a previously-known individual or by a potentially new one. However, neither these approaches, nor our system, can discriminate 1) between two unknown individuals, or 2) between two different vocalisations produced by the same unknown individual.

Finally, the multi-task design makes the system easily extendable to other tasks. Any behavioural information that can be classified can be incorporated as an additional sub-task; for instance, the acoustic type of the vocalisation (Bermant et al., 2019; Cohen et al., 2022; Oikarinen et al., 2019), its behavioural context, or its emotional valence (Briefer, 2012; Briefer et al., 2022; Marler and Evans, 1996. Briefer et al. (2022) recently proposed a system that could classify the emotional state of pigs by their vocalisations. The Rookognise system should be able to process these other informations as additional sub-tasks, which could have great implications, for instance for the assessment of welfare and management of captive animals (Laurijs et al., 2021). Finally, the system could be deployed in the wild to study the behaviours of any stable social group, so long as their identities are established.

## 5 Conclusions

In this paper, we proposed the Rookognise system as the first vocalisation-independent acoustic individual identification approach applicable to raw recordings, along with a large annotated dataset of rook vocalisation. The Rookognise system is designed to identify individual animals regardless of the vocal diversity exhibited at any level from the individual to the species. We leveraged several different mechanisms to help ensure good robustness for the problem at hand of automatic acoustic individual identification as well as great flexibility for the system to be deployed in other cases, including with additional or different tasks. The system leverages multi-task learning to improve generalisation performance on unseen data. This has two other consequences for users: first, the multiple outputs can allow separate interpretation and facilitate the extraction of particular vocalisations if desired. Second, the system is designed to be easily extendable to other tasks by simply adding or replacing heads corresponding to the task at hand. Our implementation also uses multi-channel audio to capture additional information to improve performance, although single-channel audio still provides acceptable performance at much lower data storage costs. Future work will focus on extending the system toward the open-set recognition task, and on studying the internal representations learned by the network; for instance, to test whether the network identifies individuals based on vocalisation-independent features or if different signatures exist for different vocalisation types, as has been suggested in zebra finches finches (Elie and Theunissen, 2018).

## 6 Acknowledgements

This project was funded by a Michelin Foundation donation to V.D. (N°349-PE-3176). We are grateful to the Eurometropole of Strasbourg for granting us the use of the terrain for housing the animals. We also thank IRCAM for granting us access to the AudioSculpt software.

## Footnotes

1 https://gitlab.com/kimartin/rook-vocalisation-detection

2 https://doi.org/10.5281/zenodo.6091940

## References

Adi, K., Johnson, M. T., & Osiejuk, T. S. (2010). Acoustic censusing using automatic vocalization classification and identity recognition. The Journal of the Acoustical Society of America, 127(2), 874–883. https://doi.org/10.1121/1.3273887

Beecher, M. D. (1989). Signalling systems for individual recognition: An information theory approach. Animal Behaviour, 38(2), 248–261. https://doi.org/10.1016/S0003-3472(89)80087-9

Benti, B., Curé, C., & Dufour, V. (2019). Individual signature in the most common and context-independent call of the Rook (Corvus frugilegus). The Wilson Journal of Ornithology, 131(2), 373. https://doi.org/10.1676/18-41

Bermant, P. C., Bronstein, M. M., Wood, R. J., Gero, S., & Gruber, D. F. (2019). Deep Machine Learning Techniques for the Detection and Classification of Sperm Whale Bioacoustics. Scientific Reports, 9(1), 1–10. https://doi.org/10.1038/s41598-019-48909-4

Blumstein, D. T., Mennill, D. J., Clemins, P., Girod, L., Yao, K., Patricelli, G., Deppe, J. L., Krakauer, A. H., Clark, C., Cortopassi, K. A., Hanser, S. F., Mccowan, B., Ali, A. M., & Kirschel, A. N. G. (2011). Acoustic monitoring in terrestrial environments using microphone arrays : applications, technological considerations and prospectus. Journal of Applied Ecology, (48), 758–767. https://doi.org/10.1111/j.1365-2664.2011.01993.x

Boeckle, M., Szipl, G., & Bugnyar, T. (2012). Who wants food? Individual characteristics in raven yells. Animal Behaviour, 84(5), 1123–1130. https://doi.org/10.1016/j.anbehav.2012.08.011

Boeckle, M., Szipl, G., & Bugnyar, T. (2018). Raven food calls indicate sender’s age and sex. Frontiers in Zoology, 15(1), 1–9. https://doi.org/10.1186/s12983-018-0255-z

Bogaards, N., Röbel, A., & Rodet, X. (2004). Sound Analysis and Processing with AudioSculpt 2. Proc. Int. Computer Music Conference (ICMC), 2–5. http://hdl.handle.net/2027/spo.bbp2372.2004.131

Boucherie, P. H., Mariette, M. M., Bret, C., & Dufour, V. (2016). Bonding beyond the pair in a monogamous bird: Impact on social structure in adult rooks (Corvus frugilegus). Behaviour, 153(8), 897–925. https://doi.org/10.1163/1568539X-00003372

Bradbury, J. W., & Vehrencamp, S. L. (1998). Principles of Animal Communication (2nd Edition). Sinauer Associates, Inc,.

Briefer, E. F. (2012). Vocal expression of emotions in mammals: Mechanisms of production and evidence. Journal of Zoology, 288(1), 1–20. https://doi.org/10.1111/j.1469-7998.2012.00920.x

Briefer, E. F., Sypherd, C. C., Linhart, P., Leliveld, L. M., Padilla de la Torre, M., Read, E. R., Guérin, C., Deiss, V., Monestier, C., Rasmussen, J. H., Špinka, M., Döpjan, S., Boissy, A., Janczak, A. M., Hillmann, E., & Tallet, C. (2022). Classification of pig calls produced from birth to slaughter according to their emotional valence and context of production. Scientific Reports, 12(1), 1–10. https://doi.org/10.1038/s41598-022-07174-8

Brown, E. D. (1985). The Role of Song and Vocal Imitation among Common Crows (Corvus brachyrhynchos). Zeitschrift för Tierpsychologie, 68(2), 115–136. https://doi.org/10.111/j.1439-0310.1985.tb00119.x

Campos, I. B., Fewster, R., Landers, T., Truskinger, A., Towsey, M., Roe, P., Lee, W., & Gaskett, A. (2022). Acoustic region workflow for efficient comparison of soundscapes under different invasive mammals’ management regimes. Ecological Informatics, 68. https://doi.org/https://doi.org/10.1016/j.ecoinf.2022.101554

Caruana, R. (1997). Multitask Learning (Doctoral dissertation). Carnegie Mellon University. https://doi.org/10.1007/978-1-4899-7687-1100322

Catchpole, C., & Slater, P. (2008). Bird song: Biological themes and variations, second edition (2nd Edition). https://doi.org/10.1017/CBO9780511754791

Cheng, J., Xie, B., Lin, C., & Ji, L. (2012). A comparative study in birds: Call-type-independent species and individual recognition using four machine-learning methods and two acoustic features. Bioacoustics, 21(2), 157–171. https://doi.org/10.1080/09524622.2012.669664

Christin, S., Hervet, É., & Lecomte, N. (2019). Applications for deep learning in ecology. Methods in Ecology and Evolution, 10(April), 1632–1644. https://doi.org/10.1111/2041-210X.13256

Clutton-Brock, T., & Sheldon, B. (2010). Individuals and populations: The role of long-term, individual-based studies of animals in ecology and evolutionary biology. Trends in ecology & evolution, 25, 562–73. https://doi.org/10.1016/j.tree.2010.08.002

Cohen, Y., Nicholson, D. A., Sanchioni, A., Mallaber, E. K., Skidanova, V., & Gardner, T. J. (2022). Automated annotation of birdsong with a neural network that segments spectrograms. eLife, 11, 1–32. https://doi.org/10.7554/eLife.63853

Conrady, C. R., Er, Ş., Attwood, C. G., Roberson, L. A., & de Vos, L. (2022). Automated detection and classification of southern african roman seabream using mask r-cnn. Ecological Informatics, 69. https://doi.org/https://doi.org/10.1016/j.ecoinf.2022.101593

Coombs, C. J. F. (1960). Observations on the Rook Corvus frugilegus in Southwest Cornwall. Ibis, 102(3), 394–419. https://doi.org/10.1111/j.1474-919X.1960.tb08417.x

Darras, K., Batáry, P., Furnas, B. J., Grass, I., Mulyani, Y. A., & Tscharntke, T. (2019). Autonomous sound recording outperforms human observation for sampling birds: a systematic map and user guide. Ecological Applications, 29(6). https://doi.org/10.1002/eap.1954

Dufourq, E., Batist, C., Foquet, R., & Durbach, I. (2022). Passive acoustic monitoring of animal populations with transfer learning. Ecological Informatics, 70. https://doi.org/https://doi.org/10.1016/j.ecoinf.2022.101688

Elie, J. E., & Theunissen, F. E. (2018). Zebra finches identify individuals using vocal signatures unique to each call type. Nature Communications, 9(1). https://doi.org/10.1038/s41467-018-06394-9

Fagerlund, S., & Härmä, A. (2005). Parametrization of inharmonic bird sounds for automatic recognition. 13th European Signal Processing Conference, EUSIPCO 2005, (June), 1039–1042.

Fanioudakis, L., & Potamitis, I. (2017). Deep networks tag the location of bird vocalisations on audio spectrograms. CoRR. http://arxiv.org/abs/1711.04347

Fawcett, T. (2006). An introduction to ROC analysis. Pattern Recognition Letters, 27(8), 861–874. https://doi.org/10.1016/j.patrec.2005.10.010

Ferreira, A. C., Silva, L. R., Renna, F., Brandl, H. B., Renoult, J. P., Farine, D. R., Covas, R., & Doutrelant, C. (2020). Deep learning-based methods for individual recognition in small birds. Methods in Ecology and Evolution, 11(9), 1072–1085. https://doi.org/10.1111/2041-210X.13436

Fletcher, N. H. (2000). A class of chaotic bird calls? The Journal of the Acoustical Society of America, 108(2), 821–826. https://doi.org/10.1121/1.429615

Folliot, A., Haupert, S., Ducrettet, M., Sébe, F., & Sueur, J. (2022). Using acoustics and artificial intelligence to monitor pollination by insects and tree use by woodpeckers. Science of the Total Environment, 838. https://doi.org/10.1016/j.scitotenv.2022.155883

Fox, E. J. S., Roberts, J. D., & Bennamoun, M. (2008). Call-independent individual identification in birds. Bioacoustics: The International Journal of Animal Sound and its Recording, 18:1(December 2012), 51–67. https://doi.org/10.1080/09524622.2008.9753590

Fristrup, K. M., & Mennitt, D. (2012). Biacoustical monitoring in terrestrial environments. Acoustics Today, 8(3), 16–24. https://doi.org/10.1121/1.4753913

Fu, X., Liu, Y., & Liu, Y. (2022). A case study of utilizing yolot based quantitative detection algorithm for marine benthos. Ecological Informatics, 70. https://doi.org/https://doi.org/10.1016/j.ecoinf.2022.101603

Goffinet, J., Brudner, S., Mooney, R., & Pearson, J. (2021). Low-dimensional learned feature spaces quantify individual and group differences in vocal repertoires. eLife, 10, 1–23. https://doi.org/10.7554/eLife.67855

Grießmann, B., & Naguib, M. (2002). Song Sharing in Neighboring and Non-Neighboring Thrush Nightingales (Luscinia luscinia) and its Implications for Communication. Ethology, 14, 377–387. https://doi.org/10.1046/j.1439-0310.2002.00781.x

Grill, T., & Schlöter, J. (2017). Two convolutional neural networks for bird detection in audio signals. 25th European Signal Processing Conference, EUSIPCO 2017, 2017-Janua, 1764–1768. https://doi.org/10.23919/EUSIPCO.2017.8081512

Hausberger, M., Richard-Yris, M., Henry, L., Lepage, L., & Schmidt, I. (1995). Song Sharing Reflects the Social Organization in a Captive Group of European Starlings (Sturnus vulgaris). Journal of Comparative Psychology, 109(3), 222–241. https://doi.org/10.1037/0735-7036.109.3.222

Ioffe, S. (2017). Batch Renormalization: Towards reducing minibatch dependence in batch-normalized models. Advances in Neural Information Processing Systems, 2017-Decem, 1946–1954.

Ioffe, S., & Szegedy, C. (2015). Batch normalization: Accelerating deep network training by reducing internal covariate shift. 32nd International Conference on Machine Learning, ICML 2015, 1, 448–456.

Jansen, D. A., Cant, M. A., & Manser, M. B. (2012). Segmental concatenation of individual signatures and context cues in banded mongoose (Mungos mungo) close calls. BMC Biology, 10(1), 97. https://doi.org/10.1186/1741-7007-10-97

Kahl, S., Stöter, F. R., Goeäu, H., Glotin, H., Planqué, R., Vellinga, W. P., & Joly, A. (2019). Overview of BIRDCLEF 2019: Large-scale bird recognition in soundscapes. CEUR Workshop Proceedings, 2380, 9–12.

Kahl, S., Wood, C. M., Eibl, M., & Klinck, H. (2021). BirdNET: A deep learning solution for avian diversity monitoring. Ecological Informatics, 61(January), 101236. https://doi.org/10.1016/j.ecoinf.2021.101236

Keenan, S., Mathevon, N., Stevens, J. M., Nicolé, F., Zuberböhler, K., Guéry, J. P., & Levréro, F. (2020). The reliability of individual vocal signature varies across the bonobo’s graded repertoire. Animal Behaviour, 169, 9–21. https://doi.org/10.1016/j.anbehav.2020.08.024

Kershenbaum, A., Blumstein, D. T., Roch, M. A., Akçay, Ç., Backus, G., Bee, M. A., Bohn, K., Cao, Y., Carter, G., Cäsar, C., Coen, M., Deruiter, S. L., Doyle, L., Edelman, S., Ferreri-Cancho, R., Freeberg, T. M., Garland, E. C., Gustison, M., Harley, H. E., … Zamora-Gutierrez, V. (2014). Acoustic sequences in non-human animals: A tutorial review and prospectus. Biological Reviews, 91(1), 000–000. https://doi.org/10.1111/brv.12160

Kershenbaum, A., Sayigh, L. S., & Janik, V. M. (2013). The Encoding of Individual Identity in Dolphin Signature Whistles: How Much Information Is Needed? PLoS ONE, 8(10), 1–7. https://doi.org/10.1371/journal.pone.0077671

Kirschel, A. N., Earl, D. A., Yao, Y., Escobar, I. A., Vilches, E., Vallejo, E. E., & Taylor, C. E. (2009). Using songs to identify individual mexican antthrush Formicarius moniliger: Comparison of four classification methods. Bioacoustics, 19(1-2), 1–20. https://doi.org/10.1080/09524622.2009.9753612

Kondo, N., Izawa, E. I., & Watanabe, S. (2010). Perceptual mechanism for vocal individual recognition in jungle crows (Corvus macrorhynchos): Contact call signature and discrimination. Behaviour, 147(8), 1051–1072. https://doi.org/10.1163/000579510X505427

Kong, Q., Xu, Y., & Plumbley, M. D. (2017). Joint detection and classification convolutional neural network on weakly labelled bird audio detection. 25th European Signal Processing Conference, EUSIPCO 2017, 2017-Janua, 1749–1753. https://doi.org/10.23919/EUSIPCO.2017.8081509

Laiolo, P., Palestrini, C., & Rolando, A. (2000). A study of Choughs’ vocal repertoire: Variability related to individuals, sexes and ages. Journal fur Ornithologie, 141(2), 168–179. https://doi.org/10.1046/j.1439-0361.2000.00074.x

Laurijs, K. A., Briefer, E. F., Reimert, I., & Webb, L. E. (2021). Vocalisations in farm animals: A step towards positive welfare assessment. Applied Animal Behaviour Science, 236, 105264. https://doi.org/10.1016/j.applanim.2021.105264

Lemasson, A., Ouattara, K., Petit, E. J., & Zuberböhler, K. (2011). Social learning of vocal structure in a nonhuman primate ? BMC Evolutionary Biology, (362).

Li, W., Zheng, T., Yang, Z., Li, M., Sun, C., & Yang, X. (2021). Classification and detection of insects from field images using deep learning for smart pest management: A systematic review. Ecological Informatics, 66. https://doi.org/https://doi.org/10.1016/j.ecoinf.2021.101460

Liaqat, S., Bozorg, N., Jose, N., Conrey, P., Tamasi, A., & Johnson, M. T. (2018). Domain Tuning Methods For Bird Audio Detection. https://github.com/UKYSpeechLab/ukybirddet

Liebel, L., & Körner, M. (2018). Auxiliary Tasks in Multi-task Learning, 1–8. https://doi.org/10.48550/arXiv.1805.06334

Lin, T. Y., Goyal, P., Girshick, R., He, K., & Dollar, P. (2020). Focal Loss for Dense Object Detection. IEEE Transactions on Pattern Analysis and Machine Intelligence, 42(2), 318–327. https://doi.org/10.1109/TPAMI.2018.2858826

Linhart, P., Osiejuk, T. S., Budka, M. Šálek, M., Špinka, M., Policht, R., Syrová, M., & Blumstein, D. T. (2019). Measuring individual identity information in animal signals: Overview and performance of available identity metrics. Methods in Ecology and Evolution, 10(9), 1558–1570. https://doi.org/10.1111/2041-210X.13238

Lostanlen, V., Salamon, J., Farnsworth, A., Kelling, S., & Bello, J. P. (2019). Robust sound event detection in bioacoustic sensor networks. PLOS ONE, 14(10), 1–31. https://doi.org/10.1371/journal.pone.0214168

Marler, P., & Evans, C. (1996). Bird calls: Just emotional displays or something more? Ibis, 138(1), 26–33. https://doi.org/10.1111/j.1474-919x.1996.tb04310.x

Marler, P., & Slabbekoorn, H. (Eds.). (2004). Nature’s Music: The Science of Birdsong. Elsevier. https://doi.org/10.1016/B978-0-12-473070-0.X5000-2

Marzluff, J., & Angell, T. (2005). Cultural Coevolution: How the Human Bond with Crows and Ravens Extends Theory and Raises New Questions. Journal of Ecological Anthropology, 9(1), 69–75. https://doi.org/10.5038/2162-4593.9.1.5

Mates, E. A., Tarter, R. R., Ha, J. C., Clark, A. B., & McGowan, K. J. (2015). Acoustic profiling in a complexly social species, the American crow: Caws encode information on caller sex, identity and behavioural context. Bioacoustics, 24(1), 63–80. https://doi.org/10.1080/09524622.2014.933446

McCordic, J. A., Root-Gutteridge, H., Cusano, D. A., Denes, S. L., & Parks, S. E. (2016). Calls of North Atlantic right whales Eubalaena glacialis contain information on individual identity and age class. Endangered Species Research, 30(1), 157–169. https://doi.org/10.3354/esr00735

Misra, D. (2019). Mish: A Self Regularized Non-Monotonic Activation Function. arXiv preprint 1908.08681. http://arxiv.org/abs/1908.08681

Morfi, V., & Stowell, D. (2018). Deep learning for audio event detection and tagging on low-resource datasets. Applied Sciences (Switzerland), 8(8). https://doi.org/10.3390/app8081397

Narang, S., Diamos, G., Elsen, E., Micikevicius, P., Alben, J., Garcia, D., Ginsburg, B., Houston, M., Kuchaiev, O., Venkatesh, G., & Wu, H. (2018). Mixed precision training. 6th International Conference on Learning Representations, ICLR 2018 - Conference Track Proceedings, 1–14. https://doi.org/10.48550/arXiv.1710.03740

Nolasco, I., & Stowell, D. (2022). Rank-Based Loss for Learning Hierarchical Representations. ICASSP, IEEE International Conference on Acoustics, Speech and Signal Processing - Proceedings, 2022-May, 3623–3627. https://doi.org/10.1109/ICASSP43922.2022.9746907

Ntalampiras, S., & Potamitis, I. (2021). Acoustic detection of unknown bird species and individuals. CAAI Transactions on Intelligence Technology, 6(3), 291–300. https://doi.org/10.1049/cit2.12007

Oikarinen, T., Srinivasan, K., Meisner, O., Hyman, J. B., Parmar, S., Fanucci-Kiss, A., Desimone, R., Landman, R., & Feng, G. (2019). Deep convolutional network for animal sound classification and source attribution using dual audio recordings. The Journal of the Acoustical Society of America, 145(2), 654–662. https://doi.org/10.1121/1.5087827

Pagliarini, S., Trouvain, N., Leblois, A., Hinaut, X., Pagliarini, S., Trouvain, N., Leblois, A., Hinaut, X., Applied, L.-d. G. A. N., Pagliarini, S., Trouvain, N., Leblois, A., & Hinaut, X. (2021). What does the Canary Say? Low-Dimensional GAN Applied to Birdsong. https://hal.inria.fr/hal-03244723v1

Pankajakshan, A., Bear, H. L., & Benetos, E. (2019). Polyphonic sound event and sound activity detection: A multi-task approach. arXiv, 1–5. https://doi.org/10.48550/arXiv.1907.05122

Pankajakshan, A., Thakur, A., Thapar, D., Rajan, P., & Nigam, A. (2018). All-conv net for bird activity detection: Significance of learned pooling. Proceedings of the Annual Conference of the International Speech Communication Association, INTERSPEECH, 2018-Septe, 2122–2126. https://doi.org/10.21437/Interspeech.2018-1522

Pelt, D. M., & Sethian, J. A. (2017). A mixed-scale dense convolutional neural network for image analysis. Proceedings of the National Academy of Sciences of the United States of America, 115(2), 254–259. https://doi.org/10.1073/pnas.1715832114

Potamitis, I., Ntalampiras, S., Jahn, O., & Riede, K. (2014). Automatic bird sound detection in long real-field recordings: Applications and tools. Applied Acoustics, 80, 1–9. https://doi.org/10.1016/j.apacoust.2014.01.001

Ptacek, L., MacHlica, L., Linhart, P., Jaska, P., & Muller, L. (2016). Automatic recognition of bird individuals on an open set using as-is recordings. Bioacoustics, 25(1), 55–73. https://doi.org/10.1080/09524622.2015.1089524

Røskaft, E., & Espmark, Y. (1982). Vocal communication by the rook Corvus frugilegus during the breeding season. Ornis Scandinavica, 13(1), 38–46. https://doi.org/10.2307/3675971

Ruder, S. (2017). An Overview of Multi-Task Learning in Deep Neural Networks. (May). http://arxiv.org/abs/1706.05098

Sainburg, T., Theilman, B., Thielk, M., & Gentner, T. Q. (2019). Parallels in the sequential organization of birdsong and human speech. Nature Communications, 10(1), 1–11. https://doi.org/10.1038/s41467-019-11605-y

Saito, T., & Rehmsmeier, M. (2015). The precision-recall plot is more informative than the ROC plot when evaluating binary classifiers on imbalanced datasets. PLoS ONE, 10(3), 1–21. https://doi.org/10.1371/journal.pone.0118432

Schlöter, J. (2018). Bird identification from timestamped, geotagged audio recordings. CEUR Workshop Proceedings, 2125(1).

Schneider, S., Taylor, G. W., Linquist, S., & Kremer, S. C. (2019). Past, present and future approaches using computer vision for animal re-identification from camera trap data. Methods in Ecology and Evolution, 10(4), 461–470. https://doi.org/10.1111/2041-210X.13133

Sevilla, A., & Glotin, H. (2017). Audio bird classification with inception-v4 extended with time and time-frequency attention mechanisms. CEUR Workshop Proceedings, 1866.

She, J., Zhan, W., Hong, S., Min, C., Dong, T., Huang, H., & He, Z. (2022). A method for automatic real-time detection and counting of fruit fly pests in orchards by trap bottles via convolutional neural network with attention mechanism added. Ecological Informatics, 70. https://doi.org/https://doi.org/10.1016/j.ecoinf.2022.101690

Shonfield, J., & Bayne, E. M. (2017). Autonomous recording units in avian ecological research: current use and future applications. Avian Conservation and Ecology, 12(1). https://doi.org/10.5751/ace-00974-120114

Smith, L. N. (2018). A Disciplined Approach To Neural Network Hyper-Parameters: Part 1 – Learning Rate, Batch Size, Momentum, and Weight Decay. arXiv, 1–21. https://doi.org/10.48550/arXiv.1803.09820

Srivastava, N., Hinton, G., Krizhevsky, A., Sutskever, I., & Salakhutdinov, R. (2014). Dropout: A simple way to prevent neural networks from overfitting. Journal of Machine Learning Research, 15, 1929–1958. https://doi.org/10.5555/2627435.2670313

Stowell, D. (2022). Computational bioacoustics with deep learning: a review and roadmap. PeerJ, 10. https://doi.org/10.7717/peerj.13152

Stowell, D., Morfi, V., & Gill, L. F. (2016). Individual identity in songbirds: Signal representations and metric learning for locating the information in complex corvid calls. Proceedings of the Annual Conference of the International Speech Communication Association, INTERSPEECH, 08-12-Sept, 2607–2611. https://doi.org/10.21437/Interspeech.2016-465

Stowell, D., Petrusková, T., Šálek, M., & Linhart, P. (2019a). Automatic acoustic identification of individual animals: Improving generalisation across species and recording conditions. J. R. Soc. Interface, 16(153). https://doi.org/10.1098/rsif.2018.0940

Stowell, D., Wood, M. D., Pamuła, H., Stylianou, Y., & Glotin, H. (2019b). Automatic acoustic detection of birds through deep learning: The first Bird Audio Detection challenge. Methods in Ecology and Evolution, 10(3), 368–380. https://doi.org/10.1111/2041-210X.13103

Szegedy, C., Vanhoucke, V., Ioffe, S., Shlens, J., & Wojna, Z. (2016). Rethinking the Inception Architecture for Computer Vision. Proceedings of the IEEE Computer Society Conference on Computer Vision and Pattern Recognition, 2016-Decem, 2818–2826. https://doi.org/10.1109/CVPR.2016.308

Takimoto, H., Sato, Y., Nagano, A. J., Shimizu, K. K., & Kanagawa, A. (2021). Using a two-stage convolutional neural network to rapidly identify tiny herbivorous beetles in the field. Ecological Informatics, 66. https://doi.org/https://doi.org/10.1016/j.ecoinf.2021.101466

Teixeira, D., Linke, S., Hill, R., Maron, M., & van Rensburg, B. J. (2022). Fledge or fail: Nest monitoring of endangered black-cockatoos using bioacoustics and open-source call recognition. Ecological Informatics, 69. https://doi.org/https://doi.org/10.1016/j.ecoinf.2022.101656

Terry, A. M., Peake, T. M., & McGregor, P. K. (2005). The role of vocal individuality in conservation. Frontiers in Zoology, 2, 1–16. https://doi.org/10.1186/1742-9994-2-10

Thompson, N. S., LeDoux, K., & Moody, K. (1994). A system for describing bird song units. Bioacoustics: The International Journal of Animal Sound and its Recording, 5, 267–279. https://doi.org/10.1080/09524622.1994.9753257

van Klink, R., August, T., Bas, Y., Bodesheim, P., Bonn, A., Fossøy, F., Høye, T. T., Jongejans, E., Menz, M. H., Miraldo, A., Roslin, T., Roy, H. E., Ruczyński, I., Schigel, D., Schäffler, L., Sheard, J. K., Svenningsen, C., Tschan, G. F., Wäldchen, J., … Bowler, D. E. (2022). Emerging technologies revolutionise insect ecology and monitoring. Trends in Ecology and Evolution, 20(20), 1–14. https://doi.org/10.1016/j.tree.2022.06.001

Wang, Y., Getreuer, P., Hughes, T., Lyon, R. F., & Saurous, R. A. (2017). Trainable frontend for robust and far-field keyword spotting. ICASSP, IEEE International Conference on Acoustics, Speech and Signal Processing - Proceedings, 5670–5674. https://doi.org/10.1109/ICASSP.2017.7953242

Weinstein, B. G. (2019). A computer vision for animal ecology. Journal of Animal Ecology, (87), 533–545. https://doi.org/10.1111/1365-2656.12780

Wright, L., & Demeure, N. (2021). Ranger21: a synergistic deep learning optimizer. http://arxiv.org/abs/2106.13731

Yorzinski, J. L., Vehrencamp, S. L., McGowan, K. J., & Clark, A. B. (2006). The Inflected Alarm Caw of the American Crow: Differences in Acoustic Structure Among Individuals and Sexes. The Condor, 108(3), 518–529. https://doi.org/10.1093/condor/108.3.518

