## Supplementary figures for "Rookognise: Acoustic detection and identification of individual rooks in field recordings using multi-task neural networks"

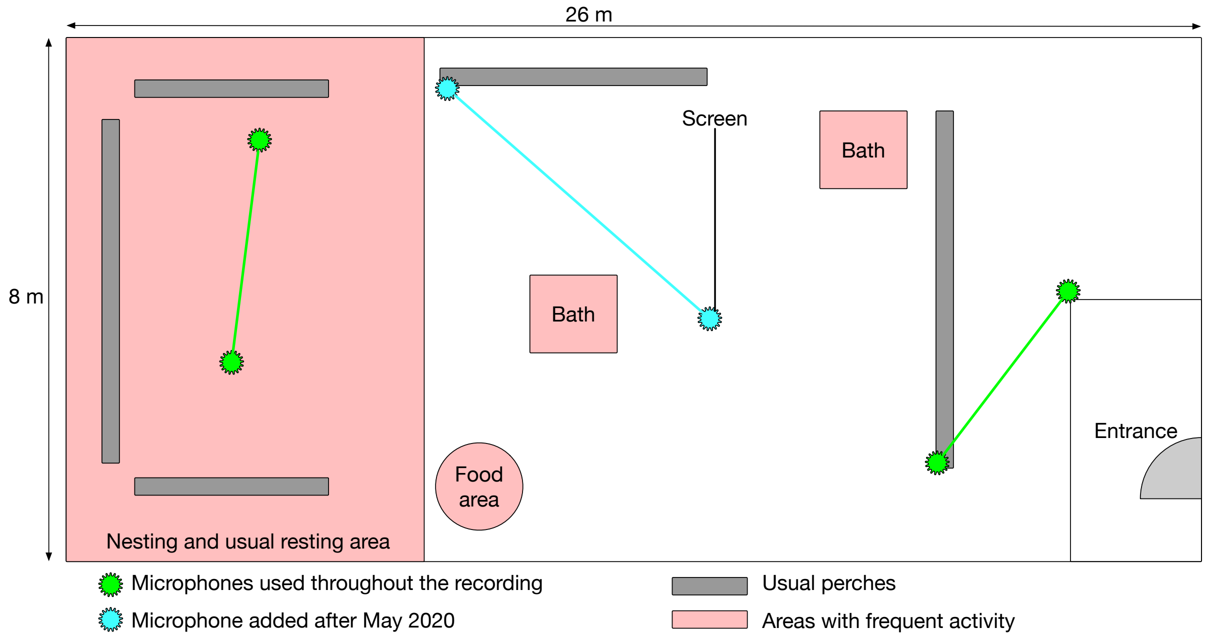


Figure S.1: Top-down schematic of the aviary, including microphone placement and areas most often used by the rooks. The aviary was entirely open with no separation between areas, except for the entrance area. Microphone location are shown as the blue and green points. Observations were interrupted between mid-March and mid-May 2020 due to the COVID lockdown. An additional microphone was placed in the middle of the aviary after observations resumed.

Table S.1: Details of the birds used in this study. The rooks were identified by coloured leg rings. All were caught from wild colonies. Different labels indicate different colonies of origin. All but H were count near Strasbourg France, while H was caught in Sountern France.

| Individual | Sex | Mate | Introduction to group | Age at introduction | Colony of origin | Note |
|---|---|---|---|---|---|---|
| A | M | G | 2013 | Unknown (juvenile) | A | |
| B | M | P, K, S[a] | 2006 | 2 months | B | |
| E | M | F, T | 2006 | 2 months | B | Sibling with J |
| F | F | E, T | 2016 | 2 months | C | Died during the recording period |
| G | F | A | 2013 | Unknown (adult) | A | |
| H | M | | 2019 | 2.5 years | Southern France | Hand-reared by an animal rescue association |
| J | F | M | 2006 | 2 months | B | Sibling with E |
| K | M | B, P | 2006 | 2 months | B | |
| L | F | O | 2016 | 2 months | C | |
| M | M | J | 2006 | 4 months | B | |
| O | M | L | 2006 | 4 months | B | |
| P | F | B, K | 2013 | Unknown (adult) | A | |
| R | F | | 2017 | 2 months | D | Died during the recording period |
| S | F | B[a] | 2013 | Unknown (adult) | A | |
| T | M | E, F | 2006 | 2 months | A | |

[a] B and S formed a pair in 2021 during the recording period. B was previously in a triadic bond with K and P for several years, and S was a single bird after losing her mate in 2019

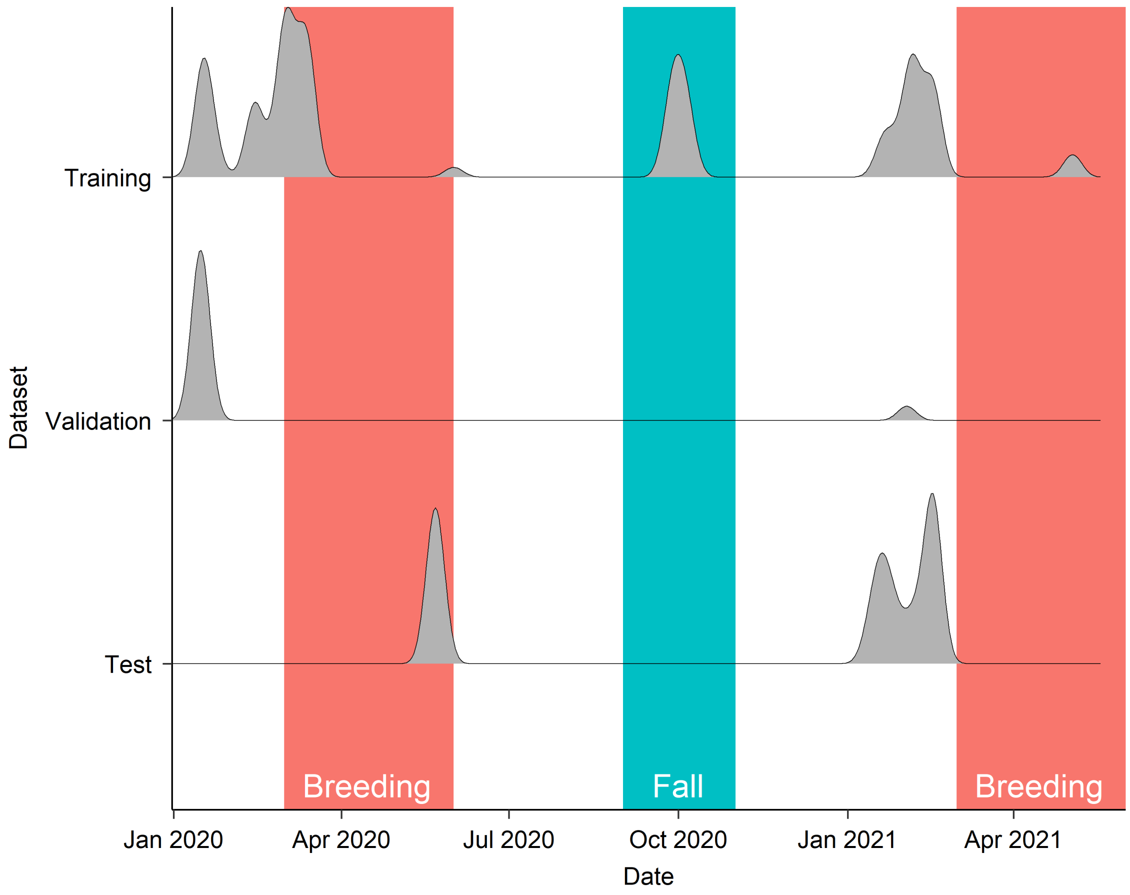


Figure S.2: Timeline of the data collection and distribution of the annotated data by period and dataset. Each line corresponds to the training, validation, or test dataset. The density lines correspond to the distribution of vocalisations over the recording period, normalized to the same maximum height. Shaded areas are periods of the year where the vocal behaviour of the rooks changed: during the breeding season (red), the males produced overall fewer vocal sequences while virtually all individuals pairs were building nests; later in the season females stayed in the nest and produced loud begging calls. During fall (blue), the same behaviours occurred at a lower intensity and without nest building.

Table S.2: The non-standard architecture designs elements included in Rookognise increase its performance. We report the effects of incorporating one of each of the non-standard architecture design elements described in Section 2.4. Each element is tested by comparing the full Rookgonise system with all the non-standard elements included, a baseline with the same architecture but all the elements replaced by their standard counterparts, and a network with the same architecture, but with one non-standard element. The training and evaluation of all the networks reported here followed the same protocol as in the main text. The baseline elements were all based on the original *sparrow* network proposed by Grill and Schluter (2017): LeakyReLU activation function with slope coefficient $\alpha = 0.01$ (replaced by Mish), Batch Normalisation (replaced by Batch Renormalisation, Renorm), Max pooling (replaced by strided convolutional pooling, Conv), Binary cross-entropy loss (replaced by the Focal loss), and Adam optimiser (replaced by Ranger21). Each respective network is denoted by Baseline + X, where X is the non-standard element included in that network. Values in bold are above the baseline, while underlined values are above the full Rookognise system. Note that the Ranger21 optimiser and the Focal loss yielded the largest improvements of the single elements. Nonetheless, the Rookognise generally performed the best in general, demonstrating that the non standard elements had non-redundant effects on performance.

| | Detection | | Sexing | | Presence | | Identification | |
|---|---|---|---|---|---|---|---|---|
| | AUROC | AUPRC | AUROC | AUPRC | AUROC | AUPRC | AUROC | AUPRC |
| Baseline | 95.90 | 90.01 | 96.80 | 84.56 | 89.49 | 67.26 | 89.93 | 60.46 |
| Baseline + Mish | 95.43 | <u>89.35</u> | 96.55 | 83.78 | <u>**89.62**</u> | 66.54 | **90.79** | **61.76** |
| Baseline + Renorm | 95.77 | 89.81 | 96.79 | 83.83 | 88.72 | <u>**68.15**</u> | **90.21** | 59.69 |
| Baseline + Conv | 95.17 | 89.01 | 96.43 | 82.05 | <u>**90.29**</u> | 62.95 | 87.95 | 57.53 |
| Baseline + Focal | <u>**97.26**</u> | <u>**91.51**</u> | **98.08** | **88.29** | 86.08 | 61.85 | **94.01** | **62.99** |
| Baseline + Ranger21 | **96.44** | <u>**90.30**</u> | **97.30** | **84.83** | <u>**90.06**</u> | <u>**70.30**</u> | **93.05** | <u>**64.73**</u> |
| Rookognise | 96.95 | 89.31 | 98.21 | 88.95 | 88.03 | 63.64 | 96.11 | 63.06 |
